# The Myogenesis Program Drives Clonal Selection and Drug Resistance in Rhabdomyosarcoma

**DOI:** 10.1101/2021.06.16.448386

**Authors:** Anand G. Patel, Xiang Chen, Xin Huang, Michael R. Clay, Natalia Komorova, Matthew J. Krasin, Alberto Pappo, Heather Tillman, Brent A. Orr, Justina McEvoy, Brittney Gordon, Kaley Blankenship, Colleen Reilly, Xin Zhou, Jackie L. Norrie, Asa Karlstrom, Jiyang Yu, Dominik Wodarz, Elizabeth Stewart, Michael A. Dyer

## Abstract

Rhabdomyosarcoma (RMS) is a pediatric cancer with features of skeletal muscle; patients with unresectable or metastatic RMS fare poorly due to high rates of disease recurrence. Here, we use single cell and single nucleus RNA-sequencing to show that RMS tumors recapitulate the spectrum of embryonal myogenesis. Using matched patient samples from a clinical trial and orthotopic patient-derived xenografts (O-PDXs), we show chemotherapy eliminates the most proliferative component with features of myoblasts; after treatment, the quiescent immature population with features of paraxial mesoderm expands to reconstitute the developmental hierarchy of the original tumor. We discovered that this paraxial mesoderm population is dependent on EGFR signaling and is sensitive to EGFR inhibitors. Taken together, this data serves as a proof-of-concept that targeting each developmental state in RMS is an effective strategy for improving outcomes by preventing disease recurrence.

## Introduction

Rhabdomyosarcoma (RMS) is the most common pediatric soft tissue sarcoma and has molecular, cellular and histopathologic features of developing skeletal muscle^1–3^. The alveolar form of RMS (ARMS) is more differentiated than the embryonal form (ERMS) and each subtype has distinct genomic and epigenomic landscapes^2,4,5^. For newly diagnosed RMS patients, the overall survival rate is 70% using multiagent chemotherapy combined with radiation and/or surgical resection^6,7^. Unfortunately, a subset of patients experience disease recurrence after treatment completion; for those patients, overall survival rate drops below 20%^8^. Genomic studies have shown that clonal selection occurs with disease recurrence, but no recurrent genetic lesion has been identified that contributes to survival of the rare clones of cells for RMS^2,5,9^. This raises the possibility that other, non-genetic mechanisms may contribute to drug resistance and disease recurrence in RMS. To explore this possibility, we performed single cell (sc) and single nucleus (sn) RNA-seq of RMS patient tumors and matched orthotopic patient-derived xenografts (O-PDXs). We also performed lentiviral barcode labeling to trace the clonal expansion of individual tumor cells during normal growth and in response to treatment. Taken together, these studies showed that individual tumor cells transition through myogenesis and the underlying myogenic developmental hierarchy contributes to clonal selection with treatment. We used the developmental program in RMS to identify therapeutic vulnerabilities that could be exploited to reduce disease recurrence. Overall, this study reveals a developmental hierarchy with RMS and introduces a novel approach to treating pediatric cancers, wherein targeting specific developmental states that are destined to persist during therapy can be used to improve treatment efficacy.

## Results

### RMS tumors have developmental heterogeneity

Skeletal muscle develops from the mesodermal cells of the somites during embryogenesis and undergoes stepwise differentiation, which is typified by the expression of myogenic regulatory factors^10,11^ (MRFs; Fig. 1A,B). RMS tumors have features of skeletal muscle including myofibers and heterogenous expression of proteins such as myogenin (MYOG)^3,12^. To further investigate the transcriptomic heterogeneity within RMS, we performed droplet-based single-cell RNA-sequencing (scRNA-seq) (Extended Data Tables 1 and 2). We obtained fresh ERMS and ARMS patient tumor tissue (Fig. 1C,D) following surgical resection and generated single-cell suspensions (>90% viable cells) for 3’-directed scRNA-seq (10x Genomics). Inference of somatic copy number alterations was used to distinguish malignant cells from non-malignant cells^13,14^ (Extended Data Fig. 1).

**Figure 1:**
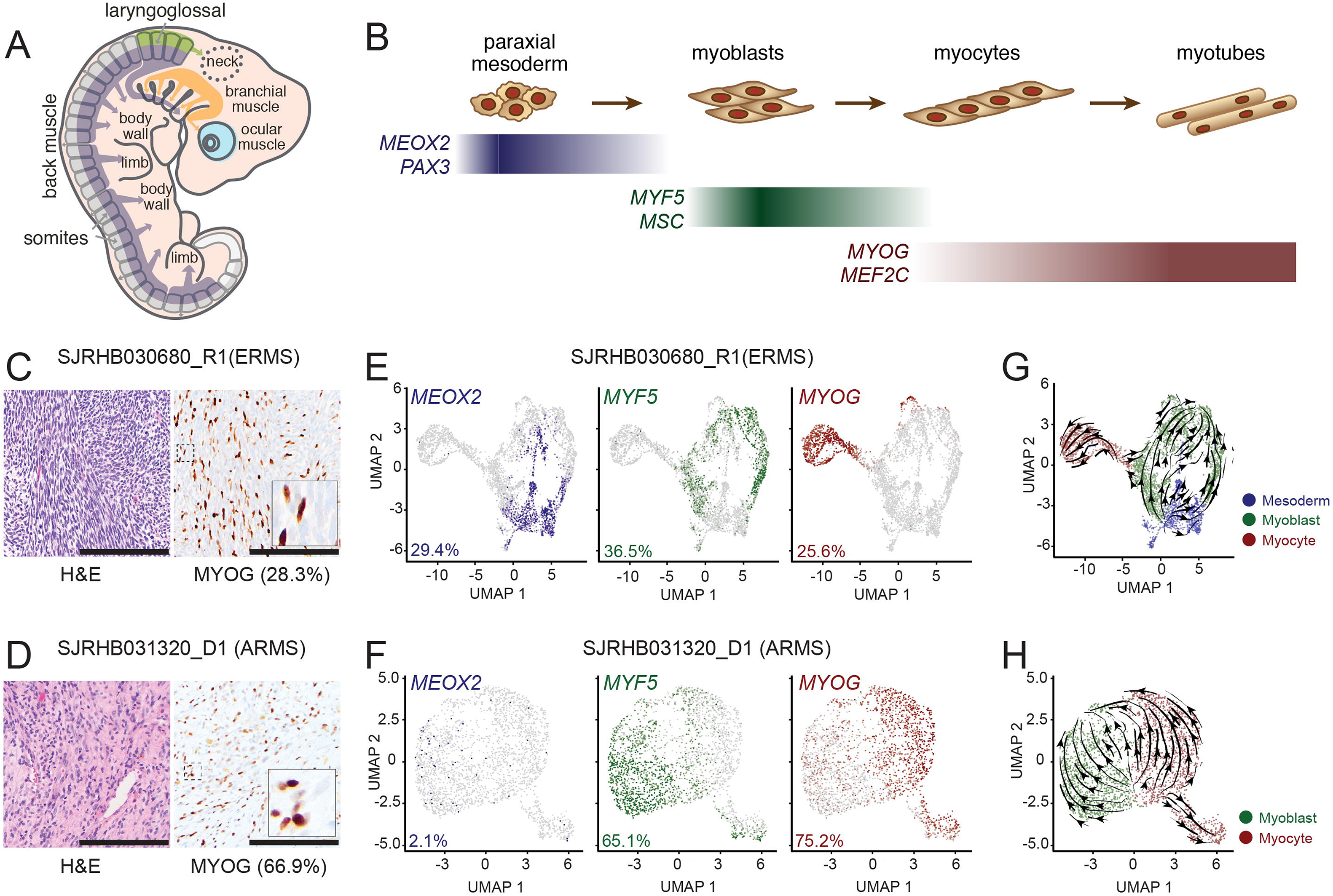
Single-cell RNA-sequencing (scRNA-seq) reveal a developmental hierarchy within RMS. **A-B,** During fetal myogenesis, mesodermal cells of the somite migrate to form skeletal muscle throughout the body (A). During that migration, these cells undergo stepwise differentiation typified by the transient expression of myogenic regulatory factors^10^ (B). **C-D,** Photomicrographs of an embryonal RMS tumor, SJRHB030680_R1 (C) and an alveolar RMS tumor, SHRHB031320_D1. *Left,* H&E staining. *Right,* Myogenin (MYOG) immunohistochemistry (IHC) with 20X magnification. *inset,* 80X magnification. **E-F,** UMAP visualization of 3,973 malignant cells from SJRHB030680_R1 (E) and 2,414 malignant cells from SJRHB031320_D1 (F). Cells are colored based on expression of *MEOX2* (*left*), *MYF5* (*center*), and *MYOG* (*right*). **G-H,** RNA velocity analysis of SJRHB030680_R1 (G) and SJRHB031320_D1 (H). Abbreviations: ERMS, embryonal rhabdomyosarcoma; UMAP, uniform manifold approximation and projection. Scale bars: C,D, 100 μm.

Single-cell analysis showed there were distinct populations of cells expressing transcription factors characteristic of paraxial mesoderm (*MEOX2, PAX3*), myoblasts (*MYF5, MSC*) and myocytes (*MYOG, MEF2C*) (Fig. 1E,F and data not shown). The proportion of *MYOG* expressing cells in the scRNA-seq data was consistent with the proportion measured by immunohistochemical staining (IHC) (Fig. 1C-F). The ARMS sample had fewer tumor cells expressing the early paraxial mesoderm MRF MEOX2 (2.1%) than the ERMS sample (29.4%), and more cells expressing the late myocyte MRF MYOG (75.2% versus 25.6%; Fig. 1E,F). RNA velocity analysis, which leverages the simultaneous measurement of spliced and unspliced RNA transcripts to generate a model of the future state of cells^15^, showed unidirectional transit of cells from the paraxial mesoderm through myoblast to the myocyte state in the ERMS tumor (Fig. 1G,H). Non-malignant cells including monocytes, fibroblasts, lymphocytes and vascular endothelial cells were readily identifiable in our scRNA-seq dataset (Extended Data Fig. 1).

The rarity of childhood cancers limits the ability to obtain fresh tissue samples for scRNA-seq. To increase the number of evaluable tumors, we validated single-nucleus RNA-sequencing (snRNA-seq) of frozen tumor tissue and adapted our computational pipeline to accommodate data generated from snRNA-seq^14^. Specifically, we compared scRNA-seq from fresh tumors (SJRHB030680_R1 and SJRHB031320_D1) (Fig. 1E,F) to snRNA-seq of matched frozen tumor specimens (Extended Data Fig. 2). As shown previously for neuroblastoma^14^, we were able to recover more cells of the tumor microenvironment (TME) by snRNA-seq compared to data generated by scRNA-seq (Extended Data Fig. 2). To extend our single cell transcriptional profiling, we performed snRNA-seq on 18 RMS tumors (12 ERMS and 6 ARMS) (Extended Data Tables 1 and 2). In total, 122,731 nuclei were analyzed from the patient tumors. As for the fresh tumors, copy number inference was used to distinguish malignant nuclei (111,474) from the normal nuclei (11,257) in the TME. The malignant nuclei were integrated using Conos, an approach that leverages inter-sample mappings to generate a unified graph for the identification of communal cell clusters^16^ (Fig. 2A). Leiden clustering identified 7 clusters, which we combined into 1 mesoderm, 4 myoblast and 2 myocyte signature groups based on expression of MRFs (Fig. 2B). The 2 myocyte populations were distinguished by expression of genes involved in muscle differentiation and function (Extended Data Table 3). The 4 myoblast populations were distinguished by ribosomal genes (p=4.3×10^−40^) and muscle differentiation genes (p=0.0005) (Extended Data Table 3). We identified 954 differentially expressed genes, of which 945 were cluster-type specific (Extended Data Table 3). Extracellular matrix and cell adhesion pathways were enriched in the paraxial mesoderm-like tumor cells, ribosome biosynthesis pathways were enriched in the myoblast-like cells and pathways involved in muscle function were enriched in the myocyte-like cells (Extended Data Table 3). While all the tumors had a mixture of cells with mesoderm, myoblast, and myocyte signatures, ARMS tumors contained significantly fewer cells with the mesodermal gene expression signature (p=0.008; unpaired t-test) and were skewed towards the myocyte signature (Fig. 2C and Extended Data Fig. 3). One ERMS tumor (SJRHB010928_R1) was notable in that it contained a majority (97%) of tumor cells with the mesodermal signature (Extended Data Fig. 3A,B). This sample was collected during treatment (Extended Data Table 1) suggesting that mesodermal cells are more resistant to treatment than the other cell populations. The proliferating cells were significantly enriched in the myoblast population (p<0.0001; one-way ANOVA with multiple comparisons) (Fig. 2E,F). All data can be viewed using an interactive viewer at: https://pecan.stjude.cloud/static/RMS-scrna-atlas-2020/.

**Figure 2:**
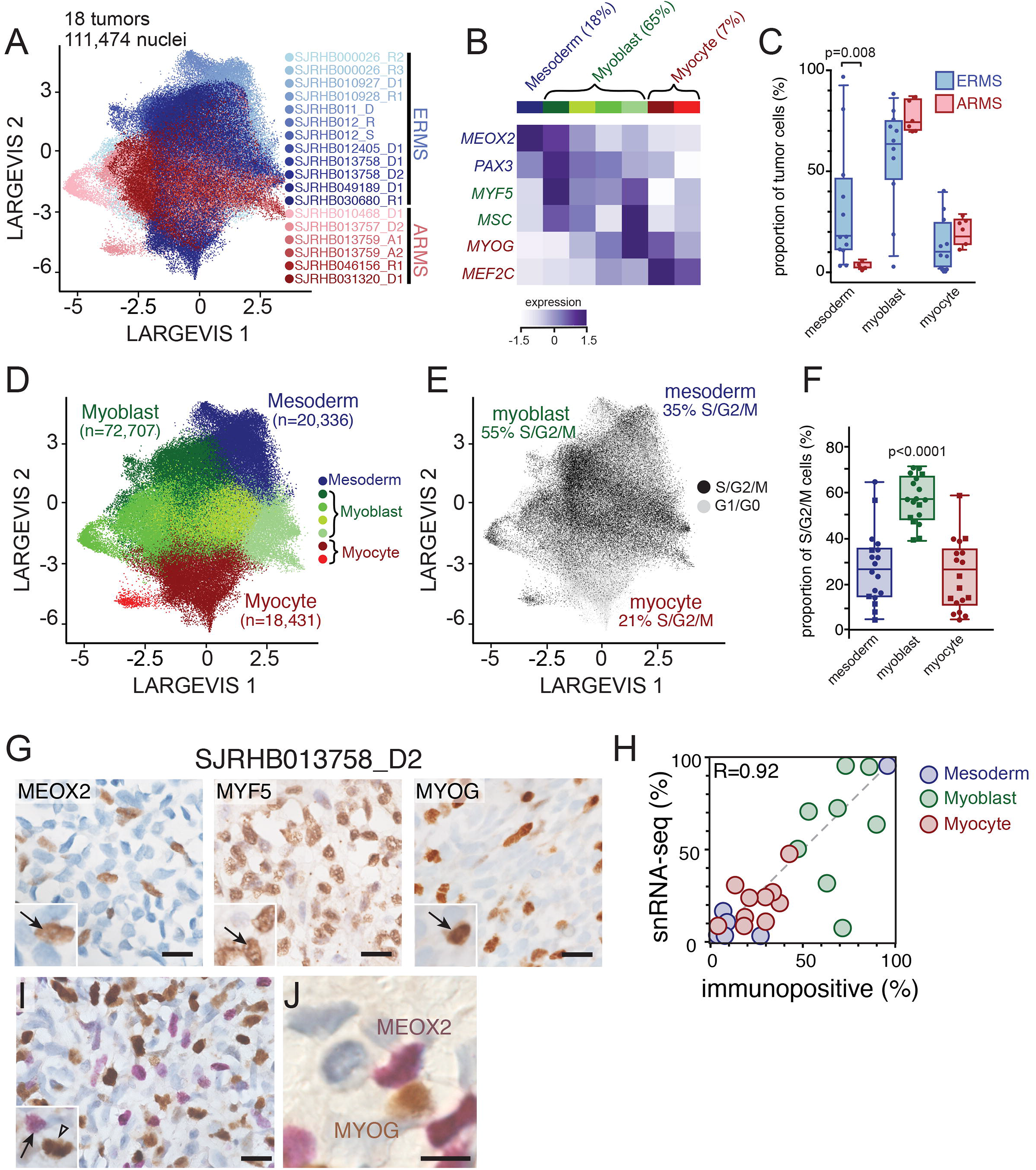
Identification of major cell clusters within patient RMS tumors using single-nucleus RNA-sequencing. **A,** LargeVis visualization of snRNA-seq of 111,474 nuclei from 18 integrated patient RMS tumors, colored based on sample. **B,** Heatmap showing expression of myogenic regulatory factor expression across seven Leiden clusters. **C,** Boxplot showing the percentage of malignant nuclei within each muscle developmental state for each tumor. The center line demarcates the median value with rectangle showing interquartile range (IQR) between the first and third quartiles. The vertical bars extending from the rectangles indicate maximum and minimum values with the exception of outliers that exceed more than 3 times the IQR. **D-E,** LargeVis visualization of Leiden clustering of snRNA-seq grouped based on expression of mesoderm, myoblast, or myocyte myogenic regulatory factors (D) or colored by predicted cell cycle phase (E). **F,** Plot of the proportion of proliferating cells (S/G2/M phase) in each group, estimated using gene signatures associated with G1, S, and G2/M phases^13^. Circles are ERMS and squares are ARMS. Center line and rectangle indicate the median and IQR as in panel (C). Vertical bars indicate the maximum and minimum values with the exception of outliers that exceed more than 3 times the IQR. **G,** Immunohistochemistry image of an ERMS tumor, SJRHB013758_D2 stained with antibodies against MEOX2 (left), MYF5 (center) and MYOG (right). **H,** Quantitation of the percentage of cells positive for MEOX2 (blue), MYF5 (green), or MYOG (red) immunohistochemical staining (*x* axis) compared to percentage of cells within each developmental state as determined by snRNA-seq (*y* axis). **I-J,** Dual staining of MEOX2 (purple) and MYOG (brown) within SJRHB013758 (I) with magnified view (J). Abbreviations: ERMS, embryonal rhabdomyosarcoma; ARMS, alveolar rhabdomyosarcoma. Scale bars: G, 10 μm.

The same approach was used to cluster the non-malignant cells within the TME (Extended Data Figure 3C-F). Comparing normal cell populations between ERMS and ARMS showed that fibroblasts in ARMS were significantly enriched in pathways involved in extracellular matrix synthesis and organization as well as cell adhesion. In addition, *SFRP2* and *SFRP4* were significantly (p<1×10^−90^) enriched in fibroblasts from ARMS (45% and 56% of cells, respectively) relative to ERMS (1% and 3%, respectively) (Extended Data Table 4). A previous pan-cancer analysis showed that *SFRP2* and *SFRP4* represent a tightly regulated transcriptional program in cancer stroma that correlates with poor prognosis, EMT and angiogenesis across multiple cancers^17^. The *HLA-A,B,C,E* and *B2M* and *CD74* genes were significantly upregulated in lymphocytes from ARMS and *HLA-DRA, DRB1* and *DPB1* were significantly upregulated in monocytes from ERMS (Extended Data Table 4).

We next investigated the spatial heterogeneity of malignant subpopulations using single and multiplex immunohistochemistry (IHC) on 12 patient tumor specimens. Consistent with our transcriptomic findings, there was heterogenous expression of MEOX2, MYF5 and MYOG protein (Fig. 2G). The proportion of immunopositive cells were correlated with the proportion of each population from the sc/snRNA-seq (Fig. 2H). Double IHC showed that these proteins were expressed in a mutually exclusive pattern consistent with the distinct clusters of mesoderm, myoblast and myocyte populations in RMS tumors from sc/snRNA-seq (Fig. 2I,J and data not shown).

### Developmental indexing of RMS using embryonic snRNA-seq data

To extend our analysis of the developmental trajectory of RMS beyond MRFs, we analyzed our RMS data within the context of early muscle development using a single-nucleus atlas of organogenesis from mouse embryos at E9.5, E10.5, E11.5, E12.5, and E13.5^18^. We extracted data from the skeletal muscle lineage and performed trajectory analysis on half of the data to generate a training dataset (Fig. 3A-D). We then adapted latent cellular analysis (LCA)^19^ to calculate the similarity in the latent cellular space between cells in the remaining half of the skeletal muscle dataset to cells used for training; a normalized muscle developmental index was then calculated for each nucleus within the validation dataset (Fig. 3E,F). The developmental index increased with embryonic age as expected within the validation dataset (Fig. 3E,F).

**Figure 3:**
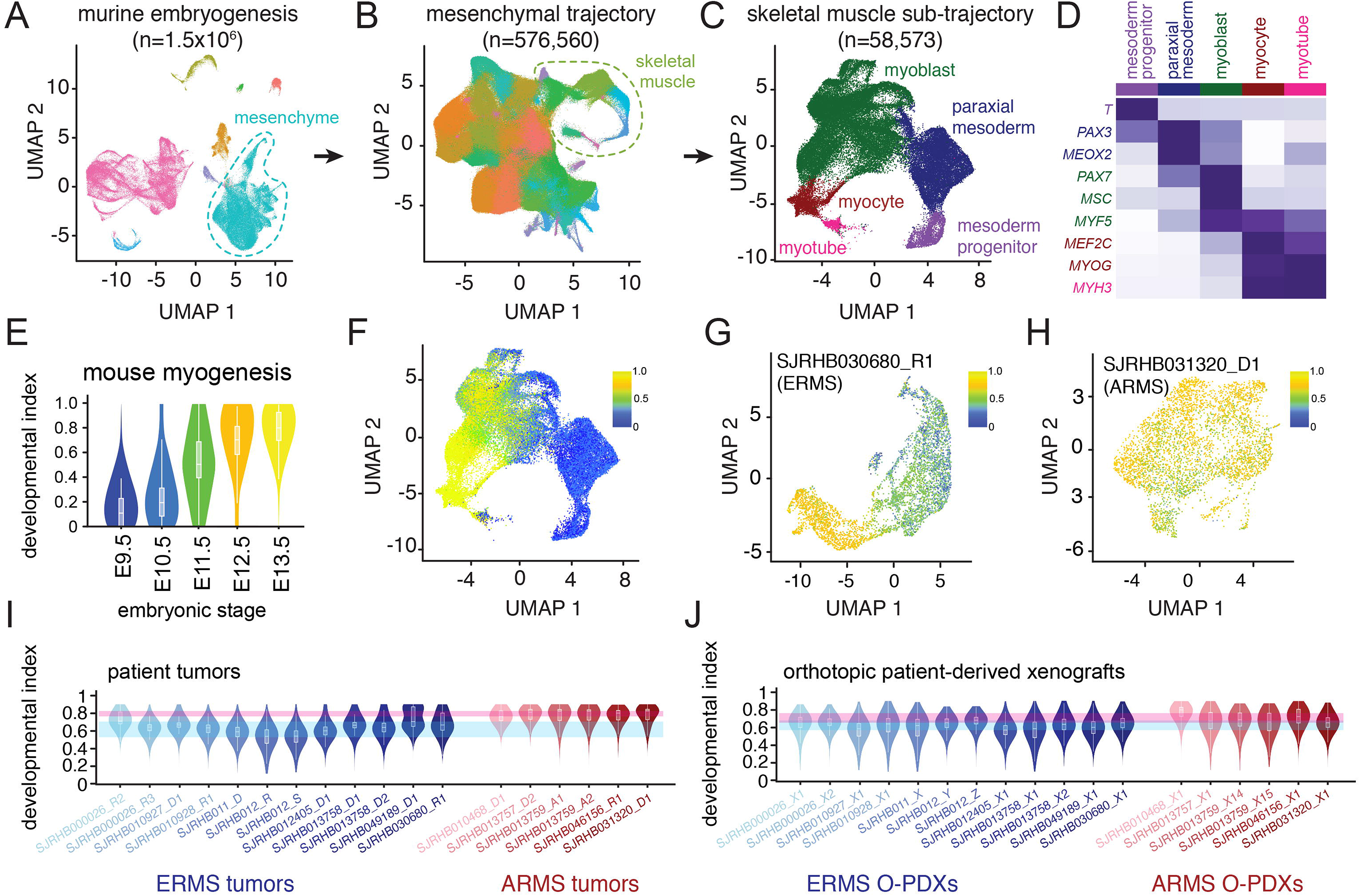
Developmental indexing of patient RMS tumors and orthotopic patient-derived xenografts. **A,** UMAP plot of 1.5 million nuclei from the Mouse Organogenesis Cell Atlas^18^, downsampled to 100,000 nuclei. Clusters are colored based on trajectory. **B,** UMAP plot of 576,560 nuclei from the mesenchymal trajectory with identification of the skeletal myogenesis sub-trajectory. Nuclei are colored based on Leiden cluster. **C,** UMAP plot of 58,573 nuclei of the skeletal muscle sub-trajectory with computational clustering that identifies nuclei from early mesodermal progenitors, paraxial mesoderm, myoblasts, myocytes and myotubes. **D,** Heatmap of aggregated transcription from each cluster demonstrating expression of myogenic regulatory factors and additional mesodermal markers. **E,** Violin plot of projected developmental indices of embryonic skeletal muscle data separated by mouse embryonic stage. **F,** UMAP plot of developmental indices within the embryonic skeletal muscle sub-trajectory. **G-H,** Application of developmental indices to an ERMS tumor, SJRHB030680_R1 (G) and an ARMS tumor, SJRHB031320_D1 (H). **I-J,** Developmental indices of 18 patient RMS tumors (I) or 18 O-PDXs (J). Abbreviations: ERMS, embryonal rhabdomyosarcoma; ARMS, alveolar rhabdomyosarcoma; UMAP, uniform manifold approximation and projection.

Using this unsupervised developmental indexing approach, we confirmed that individual RMS tumors have cellular heterogeneity that reflects normal myogenesis. For example, in SJRHB030680_R1, an ERMS tumor, we identified a broad range of developmental indices within the malignant components of the tumor (Fig. 3G). In contrast, in SJRHB031320_D1, an ARMS tumor, the range of developmental indices was narrower and more skewed toward later stages of myogenesis (Fig. 3H). Using our entire patient cohort of 18 tumors, we were able to generalize these findings to RMS tumors - ERMS tumors had a wide diversity of developmental indices while ARMS tumors narrowly centered with developmental indices from later stages of murine myogenesis (Fig. 3I).

### O-PDXs and organoids recapitulate clonal heterogeneity in RMS

We have previously established a panel of RMS O-PDXs and shared those models through the Childhood Solid Tumor Network^9^. These O-PDXs encompass the clinical and molecular diversity of RMS, and have previously undergone bulk genomic, transcriptomic, proteomic and epigenomic analyses^2,4,9^. We expanded our single-cell transcriptomic profiling to include the O-PDXs that correspond to the 18 patient tumors profiled here (Extended Data Table 2 and https://pecan.stjude.cloud/static/RMS-scrna-atlas-2020/). We performed the same analyses, including developmental indexing (Fig. 3J). All 3 cell types (mesoderm, myoblast, and myocyte) were preserved in the O-PDXs in the snRNA-seq and IHC analysis (Extended Data Fig. 4 and data not shown). As expected, the O-PDXs lacked normal cells from the patient TME but contained infiltration of murine monocytes (Extended Data Fig. 4). The patient tumor that was collected during treatment and was enriched in cells with the mesodermal signature (SJRHB010928_R1) re-established the developmental hierarchy in the O-PDX (SJRHB010928_X1) (Extended Data Fig. 4).

To complement the O-PDXs, we also evaluated the transcriptomic heterogeneity of ex vivo organoids derived from the O-PDXs (Supplemental Methods). Malignant cells within organoids shared the cellular diversity seen in the originating patient tumor and O-PDX by single cell transcriptional profiling (Extended Data Fig. 4). IHC for MEOX2, MYF5 and MYOG for the organoids showed a similarity to their matched patient tumor and corresponding O-PDX (Extended Data Fig. 4 and data not shown).

### RMS cells transition through developmental states

RNA velocity analysis (see Fig. 1G,H) suggests that individual RMS tumor cells may transition through developmental stages from mesoderm to myoblast and myocyte (Fig. 4A). Alternatively, it is possible that there are distinct clones of cells that are restricted to their developmental stage (Fig. 4B). To distinguish between these two possibilities, we used a lentiviral barcoding library that incorporates a unique oligonucleotide barcode into the 3’-untranslated region of blue fluorescent protein (BFP)^20,21^ (Fig. 4C,D). We infected 15 of the O-PDX models with the barcode library and analyzed the barcode distribution in vivo by scRNA-seq. Following scRNA-seq library generation, the barcode is retrievable by a separate PCR amplification step. In each of the tumors that we analyzed, individual barcodes were found across all tumor cell types (mesoderm, myoblast and myocytes) (Fig. 4E-G and Extended Data Table 5). Taken together, these lineage tracing data, RNA-velocity analyses and genetic clonal analysis are consistent with a model in which individual ERMS tumor cells can transition through developmental stages. The same was true for ARMS tumors but the population of cells with paraxial mesoderm gene expression signature was lower so some barcodes were found only in the myoblast and myocyte population (Extended Data Table 5).

**Figure 4:**
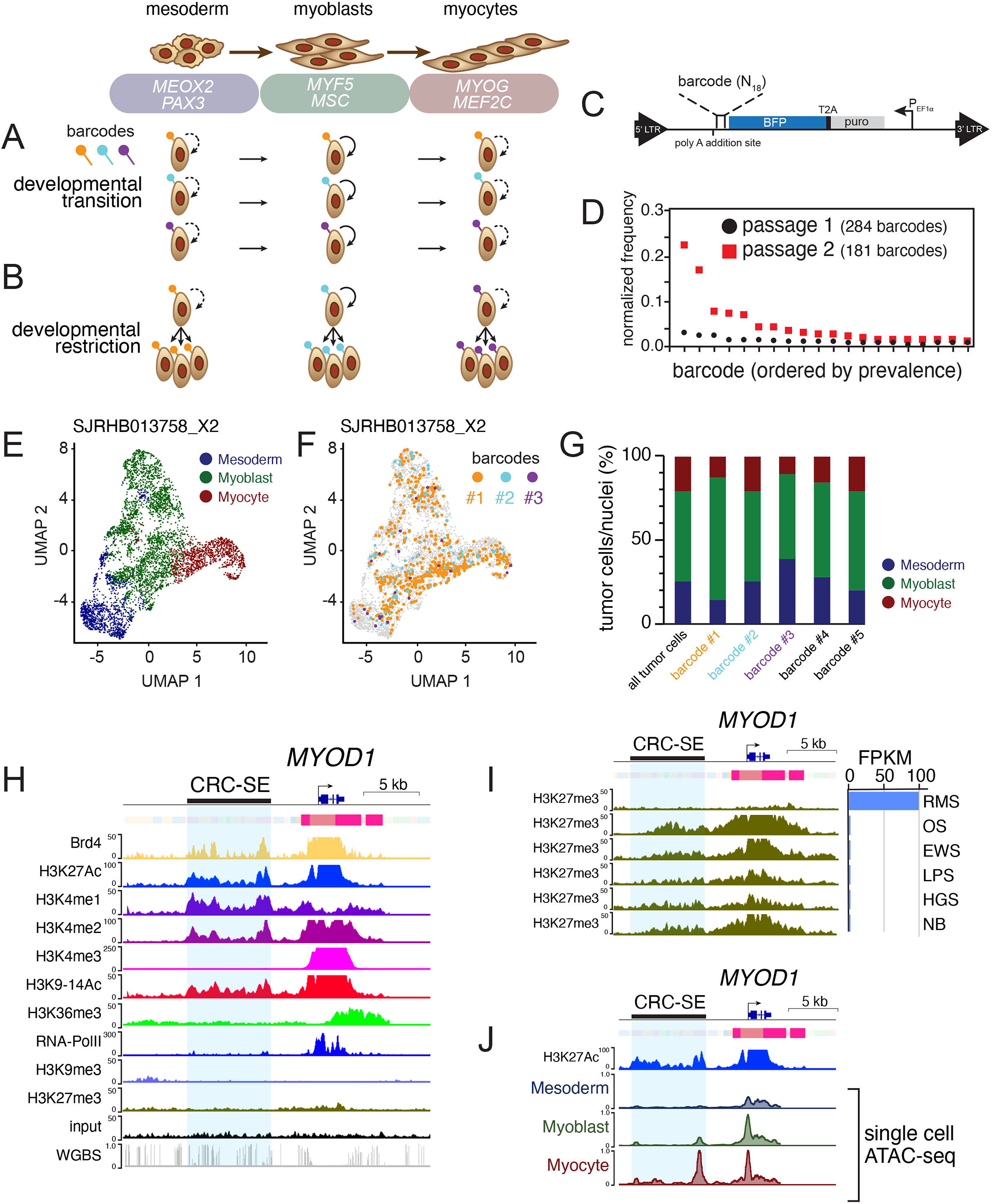
Developmental status in ERMS is plastic and associated with chromatin accessibility at core regulatory superenhancer regions. **A-B,** Two competing models of tumor heterogeneity within RMS. In the first model, RMS cells transition across developmental states (A); in the alternate model, genetically distinct clones are restricted to muscle developmental states (B). **C,** Schematic of the lentiviral barcode plasmid.^20,21^ An 18-mer of random nucleotides is incorporated into the 3’-untranslated region of a blue fluorescent protein (BFP) tag, enabling barcode recovery from scRNA-seq libraries. **D,** Plot of frequency of individual barcodes for subsequent passages of an individual ERMS O-PDX, SJRHB00026_X1. **E-F.** UMAP plot of an ERMS O-PDX SJRHB013758_X2, colored based on developmental stage (E), or with 3 specific barcodes highlighted (F). **G,** Quantitation of the developmental state diversity of all tumor cells within SJRHB013758_X2, and from the 5 most prevalent barcoded clones. **H,** ChIP-seq and chromHMM of *MYOD1* in an ERMS O-PDX, SJRHB10927_X1. Scales are indicated on the left, and a previously identified CRC-SE^4^ is highlighted in blue. **I,** Comparison of H3K27 trimethylation in various pediatric O-PDXs. OS, osteosarcoma; EWS, Ewing sarcoma; LPS, liposarcoma; HGS, high-grade sarcoma; NB, neuroblastoma. **J,** Single-cell ATAC-seq of SJRHB010927_X1 at the *MYOD1* locus; cell identities were defined via gene activity estimation, and dataset integration with scRNA-seq data^30^. Abbreviations: ERMS, embryonal rhabdomyosarcoma; ARMS, alveolar rhabdomyosarcoma; UMAP, uniform manifold approximation and projection; RMS, rhabdomyosarcoma; OS, osteosarcoma; EWS, Ewing sarcoma; LPS, liposarcoma; HGS, high grade sarcoma; NB, neuroblastoma.

### Tumor cell heterogeneity reflects differential enhancer activity

Several of the MRF genes that are turned on and off as cells transition through developmental stages have core regulatory circuit super-enhancers^4^ (CRC-SEs) (Fig. 4H and Extended Data Table 6). For example, *MEOX2* and *NFIX* (mesoderm), *PAX7* and *CREB5* (myoblast) and *FOXO1* and *SOX6* (myocyte) each have CRC-SEs (Extended Data Table 6). To determine if the chromatin accessibility of those CRC-SEs changes as individual cells transition through the myogenic differentiation program, we performed droplet-based single-cell assay of transposase-accessible chromatin sequencing (scATAC-seq) on 7 O-PDX tumors. We integrated scATAC-seq and scRNA-seq profiles to investigate the chromatin accessibility of CRC-SEs for MRFs in developmentally distinct subpopulations (Fig. 4H-J and Extended Data Table 6). Transferring cell labels between scRNA-seq data and scATAC-seq data in SJRHB010927_X1 enabled us to identify cell-type specific enhancer regions in *MYOD1*, *MSC*, *MEOX2* and several other myogenic genes (Fig. 4H-J and Extended Data Table 6). These regions correspond to previously reported core regulatory circuit domains for those genes^4,22^. Analysis of all 7 O-PDX tumors showed CRC-SEs that change in their chromatin accessibility in tumor cells with mesoderm (*MEOX2, SMAD3*), myoblast (*CREB5, PAX7*), and myocyte (*MYOD1, FOXO1*) features (Extended Data Table 6 and Extended Data Fig. 5). Collectively, these scATAC-seq studies indicate that heterogeneity of developmental states within RMS tumors is reflected in chromatin dynamics for myogenic CRC-SEs and genes.

### The mesoderm-like RMS cells are drug resistant

Current chemotherapeutic regimens for RMS include drugs that target proliferating cells. The myoblast population has the highest proportion of dividing cells in the patient tumors, the O-PDXs, and the ex vivo organoids (Fig. 5A,B and data not shown). In a pair of matched ERMS samples obtained before and during treatment, SJRHB000026_R2 and SJRHB000026_R3 (Extended Data Fig. 3A), we noted that the post-treatment sample was skewed towards mesoderm signature-expressing cells (28.6% post-treatment versus 3.4% pre-treatment) with a concomitant reduction in cells expressing the myocyte signature (1.4% post-treatment versus 31.4% pre-treatment). Additionally, one ERMS patient tumor, SJRHB010928_R1, was obtained during treatment with fewer than 5% viable cells by histology; in this sample, the majority (96.8%) of remaining viable cells expressed the mesoderm signature (Extended Data Fig. 3). Taken together, these data suggest that the myoblast population may be more sensitive to chemotherapy and the mesoderm-like population is more likely to survive treatment. To investigate this trend further, we evaluated matched formalin-fixed paraffin embedded (FFPE) tissue from 11 patients obtained at diagnosis and mid-treatment on a single therapeutic clinical trial, RMS13 (NCT01871766). We quantitated the number of cells in each sample expressing MEOX2 and MYOG (Extended Data Table 7). There was a significant enrichment in MEOX2 immunopositive cells in the post-treatment tumors relative to the matched pre-treatment RMS samples and a corresponding decrease in MYOG immunopositive cells (Fig. 5C).

**Figure 5.**
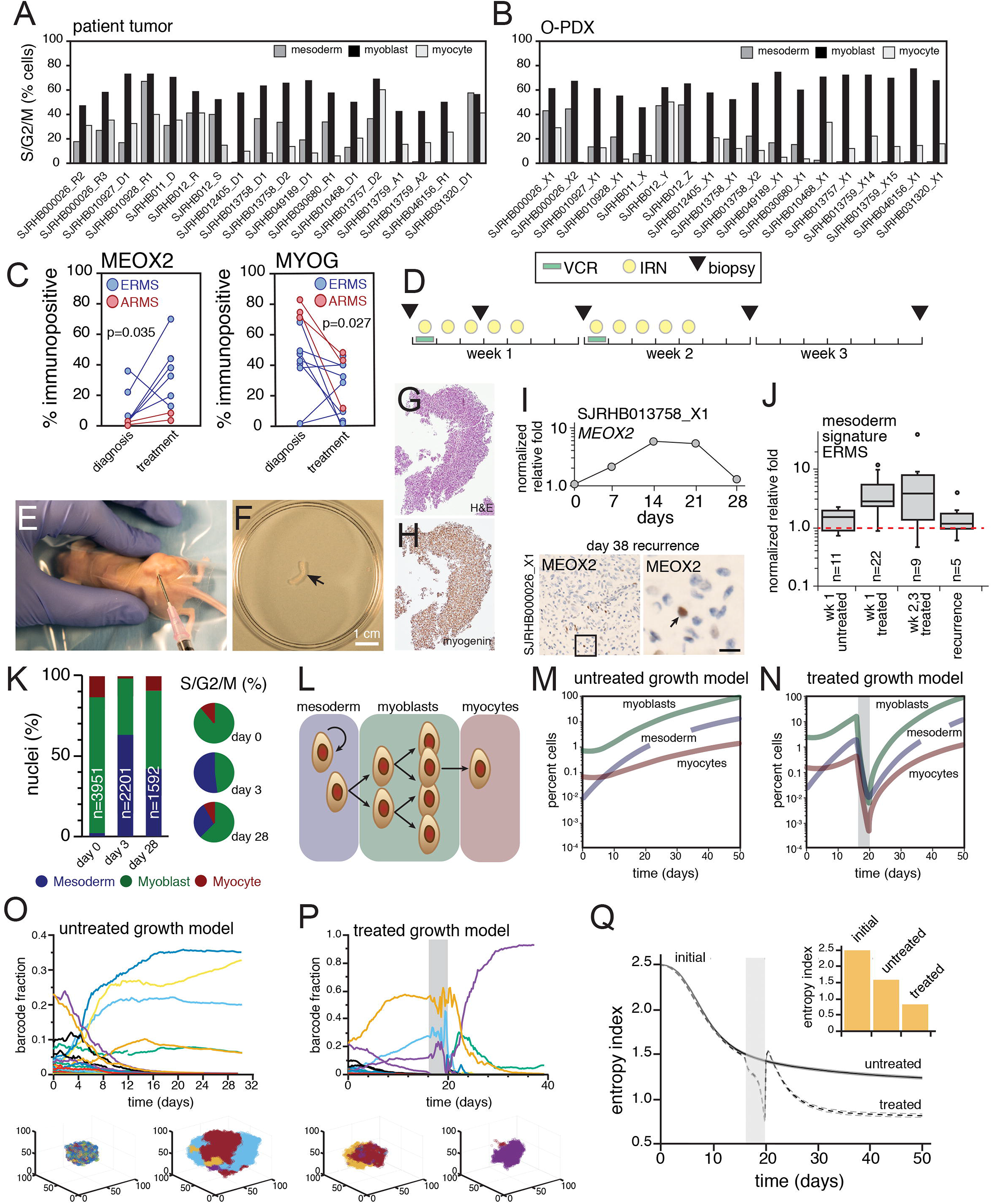
Chemotherapy treatment of ERMS selects for mesoderm developmental stages. **A-B,** Bar plots showing percentage of cells predicted to be dividing within each developmental stage for patient tumors (a) and O-PDXs (b). **C,** Plots showing immunopositivity for MEOX2 (left) and MYOG (right) in patient samples from RMS13 obtained before treatment (“diagnosis”) and during therapy (“mid-treatment”). **D,** Treatment schema for VI therapy of mice bearing RMS O-PDXs. Needle biopsies were performed at days 0, 3, 7, 14, and 21 or when tumors were large enough to sample. **E,** Photograph of needle biopsy of an orthotopically-injected xenograft. **F-H,** Photograph of tissue obtained by a biopsied O-PDX (F), which was fixed and stained using H&E (G) or MYOG (H). **I,** Plot showing longitudinal expression of *MEOX2* by qRT-PCR during treatment. There is an increase in *MEOX2* during chemotherapy (days 7,14,21) but the proportion resets to basal levels after 28 days. This was verified by IHC (lower panel). **J,** Boxplot of all biopsies for ERMS tumor bearing mice for the untreated and treated samples. The plot is an integration of expression of 6 genes (*MEOX2, PAX3, EGFR, CD44, DCN, POSTN*) expressed as normalized relative fold. The center line demarcates the median value with rectangle showing interquartile range (IQR) between the first and third quartiles. The vertical bars extending from the rectangles indicate maximum and minimum values with the exception of outliers that more than 3 times the IQR. **K,** Relative proportion of nuclei in each developmental state for longitudinal biopsies of a single O-PDX, determined using snRNA-seq of biopsied tissue. **L,** Diagram of the mathematical model of ERMS developmental heterogeneity. **M-N,** Simulated average population size for an untreated ERMS tumor (M) or a treated ERMS tumor (N) briefly exposed to an antiproliferative agent (gray bar). Average population size over 524 simulations are shown, standard error bars are too small to see. **O-P,** Simulated time course of barcode dynamics for an ERMS tumor that was either untreated (O) or briefly treated (P; duration of treatment in grey bar). Each curve represents a different barcoded lineage. One realization of the stochastic dynamics is shown. Insets under each graph show spatial distributions of bar codes (color coded) in myoblast cells at an early and late stage of tumor growth (O) and pre- and post-therapy (P). **Q,** Temporal development of the average entropy index (measure of barcode diversity) during barcoded ERMS tumor growth, either untreated or briefly treated (grey bar). Average entropy values over 524 simulations ± standard errors (dashed lines) are shown. *Inset,* bar plot comparing the initial entropy index to the final entropy index of untreated or treated tumors in experiments. Model parameters were: average value of L_mes_ =0.0035 (r_1_=1.5, r_2_=0.0001), L_blast_ =0.0045, P_mes_ =0.55, P_blast_ =0.49, D=0.035, α_mes_ =0.0014, α_blast_ =0.0035. The parameter units are per minute. Abbreviations: ERMS, embryonal rhabdomyosarcoma; ARMS, alveolar rhabdomyosarcoma; VCR, vincristine; IRN irinotecan.

To model clonal selection in the laboratory, we generated longitudinal samples from repeat biopsy of O-PDXs treated with a standard drug combination used to treat patients with RMS (vincristine (VCR) and irinotecan (IRN)) at clinically relevant doses and schedules^4,9^ (Fig. 5D). For each O-PDX (SJRHB000026_X1, SJRHB013758_X1, SJRHB011_X, SJRHB013757_X1 and SJRHB013759_X14), biopsies were performed at multiple timepoints (before treatment (day 0), day 3, day 7, day 14 and day 21 of the first course) when sufficient tumor was present to sample (Fig. 5E,F). We also collected tumor biopsies after the tumors recurred. A portion of each biopsy underwent formalin-fixation for IHC staining for MEOX2, MYF5 and MYOG (Fig. 5G,H and data not shown). The remaining biopsy portion was utilized for quantitative RT-PCR for 21 genes expressed in mesoderm, myoblast and myocyte-like RMS tumor cells or snRNA-seq. In total, 250 biopsies were collected and 6,480 qRT-PCR reactions were performed (Extended Data Table 8-13). As in patient samples, the myoblast and myocyte populations were sensitive to treatment and the mesoderm tumor cells population was enriched (Fig. 5I-K and Extended Data Table 8-13). Moreover, the normally quiescent mesoderm-like cells re-entered the cell cycle to initiate myogenesis (Fig. 5K).

Taken together, our data suggest that ERMS tumor cells transition through distinct states that represent progressive stages of myogenesis. These different states (paraxial mesoderm, myoblast, myocyte) have distinct proliferation properties and differential sensitivity to chemotherapy. To further refine our understanding of the cellular heterogeneity of ERMS tumors, their developmental trajectory and clonal selection with treatment, we developed a mathematical model that follows the fate of cells in both 3-dimensional space and time. Experimentally determined barcode distribution in each compartment was used to develop the model (Fig. 5L), and barcode diversity was tracked over time. We assumed that upon cell division, cells maintain their barcodes and we included barcoded and non-barcoded cells to reflect the in vivo experiments. The relative proportion of different division types (self-renewing/differentiating) in the mesodermal compartment determines whether the tissue remains in homeostasis and influences the degree of clonal diversity loss over time. To parameterize the model, we used experimental data from 10 barcoded ERMS xenografts. The fraction of dividing cells and distribution of cells across compartments was determined from the sc/snRNA-seq data. Our ERMS model predicts a decrease in clonal diversity (as measured by barcode diversity) over time and clonal selection with treatment for individual tumors (Fig. 5M-P). To test this experimentally, we performed scRNA-seq on a barcoded ERMS tumor (SJRHB000026_X1) after initial labeling and a subsequent passage in mice with and without clinically relevant chemotherapy (vincristine+irinotecan). As predicted by the three-compartment model, there was a decrease in clonal diversity over time and clonal selection with treatment (Fig. 5O-Q). Additional iterations of modeling and comparison to in vivo barcode distribution data are consistent with differential cytotoxicity across the cellular populations (mesoderm, myoblast, myocyte). In particular, a subset of the mesoderm-like cells are proliferating thereby making them sensitive to chemotherapy. Based on our model, partial elimination of the mesoderm-like population is required to account for the clonal selection we observe in vivo in mice and in patients.

### EGFR is a therapeutic vulnerability in paraxial mesoderm RMS cells

Having shown that the paraxial mesoderm RMS cells are more quiescent and drug resistant than the myoblast population, we set out to identify therapeutic vulnerabilities unique to this population using a systems biology algorithm, NetBID (data-driven Network-based Bayesian Inference of Drivers)^23,24^. NetBID, which was originally developed for bulk -omics data, was adapted to analyze snRNA-seq profiles of our panel of 18 RMS patient tumors. We first used the SJARACNe algorithm^25^ to reverse engineer cell type–specific interactomes for each of the 5 major cell types from the integrated snRNA-seq profiles (Fig. 6A). With a focus on signaling drivers, we used the cell type–specific interactomes of 2,543 genes/proteins and inferred their network activities in each nucleus using the NetBID algorithm. We then performed differential activity analysis to identify cell type–specific therapeutic vulnerabilities in the RMS tumor cells with the mesodermal signature. EGFR was significantly activated in the mesoderm population compared to myoblasts (p=4.4×10^−135^) and myocytes (p−1.8×10^−174^) and the network was rewired as cells transition through the developmental hierarchy (Fig. 6B). EGFR network activity was also significantly higher in ERMS relative to ARMS (p=5.4×10^−36^) (Fig. 6C,D). These data are consistent with previous integrated epigenetic/proteomic analyses for differential pathway activity in ERMS and ARMS^4^. In addition, previous studies have shown heterogenous expression of EGFR protein in FFPE samples of RMS^26–28^. To validate these data, we performed IHC for EGFR alone and in combination with markers of each cell population. We used the 5B7 monoclonal antibody which has been previously shown to correlate with EGFR inhibitor (EGFRi) responsiveness in lung cancer^29^. There was co-localization of EGFR with MEOX2 in 2-color IHC and EGFR was mutually exclusive with MYOG (Fig. 6E,F).

**Figure 6.**
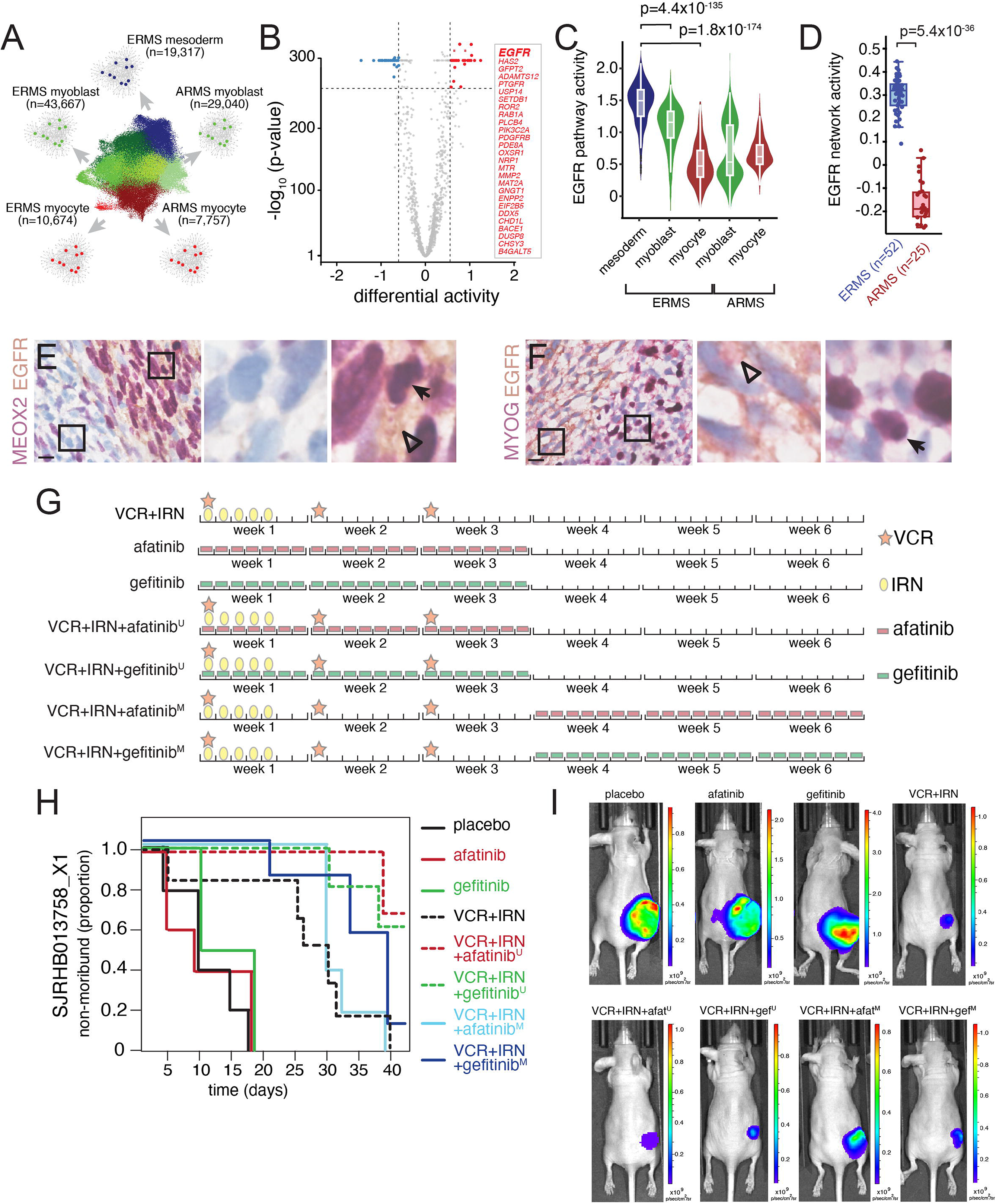
Mesoderm-like ERMS cells are uniquely vulnerable to EGFR blockade. **A,** Schematic workflow of NetBID algorithm to identify cell type–specific drivers from snRNA-seq data. **B,** Volcano plot of differential activity analysis of signaling drivers in ERMSmesoderm vs. other cell types. **C-D,** EGFR NetBID activity in different developmental states from snRNA-seq data (C) and inferred from bulk RNA-seq of patient tumors (D). **E-F,** Dual IHC staining of ERMS patient tumor, SJRHB030680_R1, combining EGFR (brown) with either MEOX2 (E) or MYOG (F) in purple. **G,** Schedules of drugs used for preclinical study. **H,** Survival curves for each treatment group for a ERMS tumor O-PDX (SJRHB01378_X1). **I,** Representative image of bioluminescence for mice treated on the study, scale bar is photons/sec/cm^2^ str. Scale bars: E,F, 10 μm. Abbreviations: VCR, vincristine; IRN, irinotecan; ERMS, embryonal rhabdomyosarcoma; ARMS, alveolar rhabdomyosarcoma.

To determine if EGFR is a therapeutic vulnerability in RMS, we exposed 3D ERMS organoids that contain all 3 cell populations (Extended Data Fig. 6) to two different EGFRi’s (gefitinib and afatinib) with increasing concentrations of SN-38, the active metabolite of irinotecan (Supplemental Information). The EGFRi’s alone had no effect on overall organoid viability as measured with CellTiter-Glo 3D which is not surprising given the low percentage of mesoderm-like cells in the organoids (Extended Data Fig. 6). However, when the proliferating myoblast population was targeted with increasing concentrations of SN-38, the addition of EGFRi’s significantly reduced viability (Extended Data Fig. 6). In a representative ERMS O-PDX (SJRHB013758_X1), there was a significant improvement in outcome when afatinib or gefitinib was combined with IRN+VCR (Fig. 6H,I). For a second ERMS O-PDX with a low percentage of mesoderm-like cells, SJRHB010927_X1, O-PDXs responded robustly to IRN+VCR therapy alone, reinforcing the importance of including standard-of-care treatment in preclinical testing (Extended Data Fig. 6).

## Discussion

In conclusion, we have discovered that RMS tumor cells can transition through different stages of myogenesis from an immature quiescent paraxial mesoderm state through a highly proliferative myoblast stage and into a more differentiated myocyte state. Not only do cells undergo changes in gene expression during these developmental transitions but super-enhancer chromatin accessibility is also dynamic. While proliferating cells can be identified in tumor cell population in patient tumors and O-PDXs, the most proliferative cells are in the myoblast stage. It is therefore not surprising that broad spectrum chemotherapeutic regimens that target rapidly dividing cells efficiently reduce tumor volume by killing the myoblast-like RMS tumor cells. The immature paraxial mesoderm-like RMS tumor cells are more quiescent and can survive therapy and then expand to repopulate the myogenic lineage found in the primary tumor. These observations are consistent with decades of clinical research showing that different combinations of broad-spectrum chemotherapy or intensification of existing regimens have failed to improve outcomes for children with RMS^6,7^. Those different treatment approaches are killing the rapidly dividing myoblast-like cells and the resistant mesoderm-like cells survive and contribute to disease recurrence. By focusing our investigation on the mesoderm-like cells, we identified a dependence on EGFR that can be exploited with EGFRi in vivo. While there were only a small number of mesoderm-like cells in ARMS tumors, we discovered a dramatic upregulation of EGFR during treatment suggesting that EGFRi’s may be useful for both types of rhabdomyosarcoma. Our study shows that treatment for RMS and possibly other pediatric solid tumors should focus on total clonal elimination rather than continuing to target just the most proliferative cell population that makes up the bulk of the tumor. Such an approach may reduce disease recurrence and improve survival and quality of life for children with solid tumors.

## Materials and Methods

## Acknowledgements

We thank A. McArthur for editing the manuscript. This work was supported by Cancer Center Support (CA21765) and grants to M.A.D. from the National Institutes of Health (EY014867, EY018599, and CA168875). The content is solely the responsibility of the authors and does not necessarily represent the official views of the National Institutes of Health. Research was also supported by American Lebanese Syrian Associated Charities. M.A.D. was also supported by the Howard Hughes Medical Institute, Alex’s Lemonade Stand, and Tully Family and Peterson Foundations. A.G.P. was supported by Alex’s Lemonade Stand, Hyundai Hope on Wheels, and Damon Runyon Cancer Research Foundations.

## Author Contributions

A.G.P, X.C. and M.A.D designed the study. A.G.P processed fresh and frozen tissue for single-cell/nucleus RNA-seq and ATAC-seq. B.G., K.B., and E.S. assisted in patient sample and xenograft tissue accrual for molecular analyses. A.G.P., X.C., X.H., and J.Y. performed computational analysis of data. M.R.C., B.A.O., and H.T. supervised immunohistochemical staining and pathology review. N.K. and D.W. performed mathematical modeling of tumors. M.J.K. and A.P. wrote and supervised the RMS13 clinical trial. A.K. manages the Childhood Solid Tumour Network. J.M. and K.B. assisted in molecular analysis. C.R. and X.Z. developed the single-cell viewer portal. K.B. and E.S. performed the preclinical testing and tumor biopsies. A.G.P. and J.L.N. developed the tumor organoid model.

## Competing Interest Statement

The authors declare no competing financial interests.

## Supplementary Information

**Supplementary Information** is available for this paper.

## Corresponding Author

Correspondence and requests for materials should be addressed to Michael Dyer at.

## Data and Source Code Availability

NetBID: https://jyyulab.github.io/NetBID

All processed single-cell and single-nucleus RNA-sequencing data are publicly accessible via an online data portal (https://pecan.stjude.cloud/static/RMS-scrna-atlas-2020).

All raw sequencing data will be uploaded and be publicly available at the time of publication through GEO accession GSE174376.

## Notes

https://pecan.stjude.cloud/static/RMS-scrna-atlas-2020/

### Competing Interest Statement

The authors have declared no competing interest.

### Summary of Updates

Supplemental Tables added.

