## Supplementary Information for "The Myogenesis Program Drives Clonal Selection and Drug Resistance in Rhabdomyosarcoma"

### Supplementary Tables

**Extended Data Table 1** – Characteristics of patient tumors and O-PDXs used for sc/snRNA-seq. Basic clinical information is provided along with stage, histologic subtype, and site of disease. For those alveolar tumor samples, we provide the detected fusion type (PAX3-FOXO1 or PAX7-FOXO1).

**Extended Data Table 2 –** Sequencing metrics for sc-/snRNA-sequencing datasets. Basic sample quality control metrics are provided, including whether the processed tissue was initially fresh or frozen. Fresh tissue was processed for single-cell RNA-seq, while frozen tissue was processed for single-nucleus RNA-seq. One tab describes metrics for patient tumors, while the second tab provides information about matched O-PDXs.

**Extended Data Table 3** – Differentially expressed genes across tumor clusters. The first tab shows genes that are differentially expressed based on developmental subgrouping (mesoderm, myoblast, or myocyte). The second tab shows data from the first tab filtered such that at least one group must have 5% of cells expressing the gene, at least one group must have mean expression of 0.05 and the differential % expressing cells must be at least 4% and the differential mean must be at least 0.04. The 3rd to 5th tabs show genes that are enriched in each developmental group, followed by pathway enrichment analysis of those groups in the 6th tab. The 7th and 8th tabs show differential expression between the four myoblast subclusters and the two myocyte subclusters.

**Extended Data Table 4 –** Differentially expressed genes across non-malignant cell populations. Each tab shows genes detected in each non-malignant subpopulation (fibroblast, vascular endothelium, lymphocytes, monocytes) divided by RMS histologic subtype.

**Extended Data Table 5 –** Sequencing metrics for lentiviral barcode experiments. Individual tabs for each O-PDX with raw data showing the number of cells and percentage of cells with each lentiviral barcode divided by developmental group.

**Extended Data Table 6 –** CRC-SE analysis and scATAC-seq. The first tab reflects expression of CRC-SE genes[4] in the snRNA-seq dataset divided by developmental state. The 2nd to 8th tabs show differentially expressed peaks within the CRC-SE for each of seven O-PDXs.

**Extended Data Table 7 –** Quantitation of IHC from matched pairs samples from patients with RMS obtained before therapy and during treatment.

**Extended Data Table 8 –** qRT-PCR analysis of biopsied tissue from ERMS O-PDX SJRHB000026_X1. Individual tabs show data for a single mouse. Tabs are labelled with mouse number and treatment group (C, control; VI 50%, vincristine + 50% irinotecan dose; VI 100%, vincristine + 100% irinotecan dose). A summary statistic on the right shows the mesoderm, myoblast and myocyte signatures for each time point.

**Extended Data Table 9 –** qRT-PCR analysis of biopsied tissue from ERMS O-PDX SJRHB013758_X1. Individual tabs show data for a single mouse. Tabs are labelled with mouse number and treatment group (C, control; VI 50%, vincristine + 50% irinotecan dose; VI 100%, vincristine + 100% irinotecan dose). A summary statistic on the right shows the mesoderm, myoblast and myocyte signatures for each time point.

**Extended Data Table 10** – qRT-PCR analysis of biopsied tissue from ERMS O-PDX SJRHB011_X. Individual tabs show data for a single mouse. Tabs are labelled with mouse number and treatment group (C, control; VI 50%, vincristine + 50% irinotecan dose; VI 100%, vincristine + 100% irinotecan dose). A summary statistic on the right shows the mesoderm, myoblast and myocyte signatures for each time point.

**Extended Data Table 11** – qRT-PCR analysis of biopsied tissue from ARMS O-PDX SJRHB013757_X2. Individual tabs show data for a single mouse. Tabs are labelled with mouse number and treatment group (C, control; VI 50%, vincristine + 50% irinotecan dose; VI 100%, vincristine + 100% irinotecan dose). A summary statistic on the right shows the mesoderm, myoblast and myocyte signatures for each time point.

**Extended Data Table 12** – qRT-PCR analysis of biopsied tissue from ERMS O-PDX SJRHB013759_X14. Individual tabs show data for a single mouse. Tabs are labelled with mouse number and treatment group (C, control; VI 50%, vincristine + 50% irinotecan dose; VI 100%, vincristine + 100% irinotecan dose). A summary statistic on the right shows the mesoderm, myoblast and myocyte signatures for each time point.

**Extended Data Table 13** – Preclinical data for mice bearing xenografts treated with vincristine+irinotecan. Each tab represents raw data for a xenograft, with each row representing a single mouse.

**Extended Data Table 14** – Preclinical data for mice bearing ERMS xenografts treated with EGFRi, vincristine+irinotecan, or a combination of EGFRi+vincristine+irinotecan. Each tab represents raw data for a xenograft, with each row representing a single mouse.

### Supplementary Figures

**Extended Data Figure 1: Validation of malignant tumor cell populations by inferring copy-number variation. A-C,** Analysis of scRNA-seq scRNA-seq data from SJRHB030680_R1, showing UMAP visualization (A), heatmap for marker gene expression (B), and inferred copy number alterations (C). **D-F,** Analysis of scRNA-seq scRNA-seq data from SJRHB031320_D1, showing UMAP visualization (D), heatmap for marker gene expression (E), and inferred copy number alterations (F). Inferred CNV plots are oriented by chromosomal position (column) for each individual cell (row). Chromosome amplifications are colored red, while deletions are colored blue. Non-malignant cells are shown on the top portion (reference) of the heatmap while malignant cells are shown on the bottom (observations).

Extended Data Figure 1

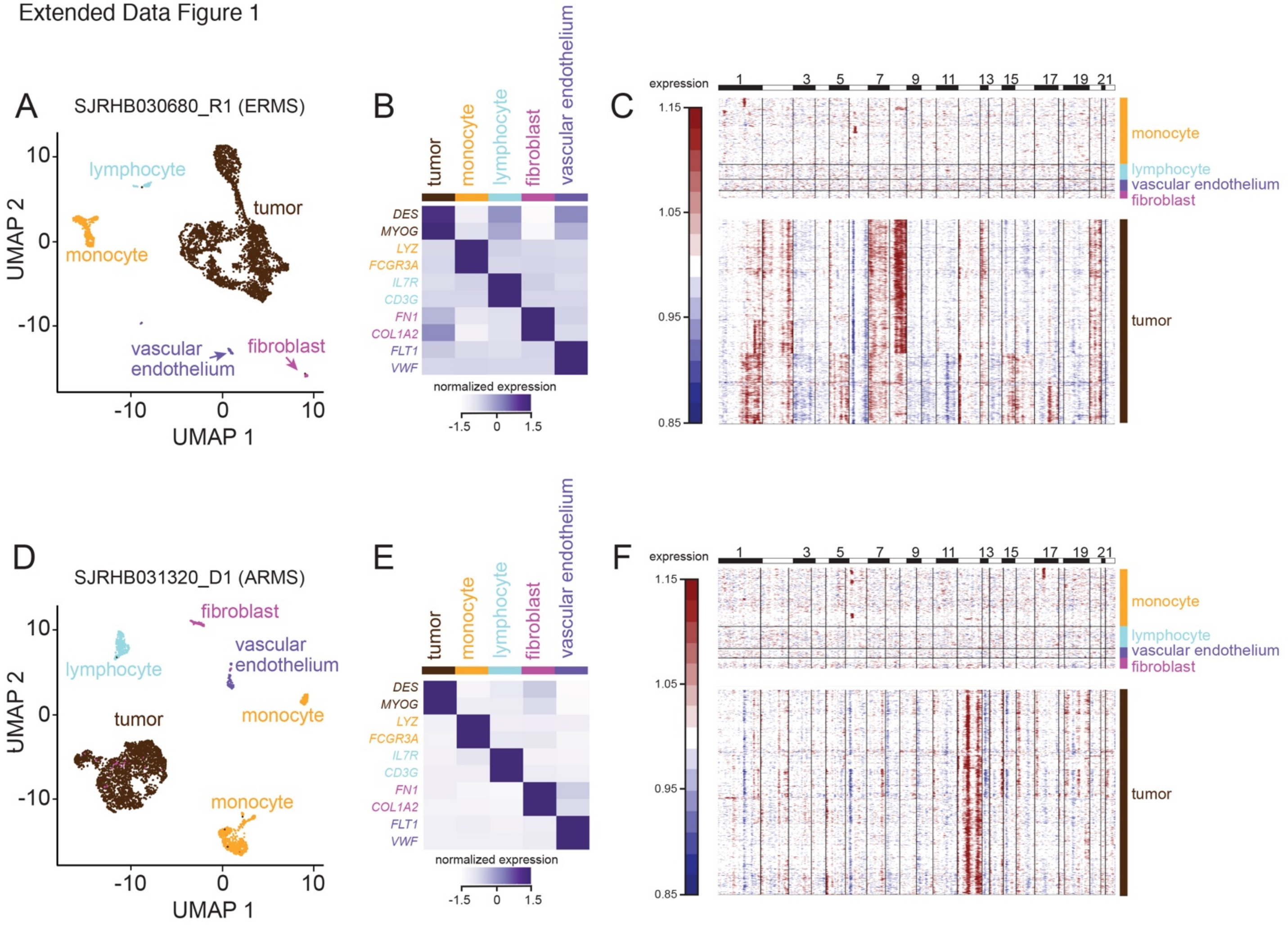

**Extended Data Figure 2: Single-nucleus RNA-sequencing preserves cell type variability. A-B,** UMAP plots comparing scRNA-seq data (A) and snRNA-seq data (B) of SJRHB030680_R1 following dataset integration. **C,** Bar graph comparing cellular diversity of single-cell and single-nucleus datasets of SJRHB030680_R1. **D-E,** UMAP plots comparing scRNA-seq data (D) and snRNA-seq data (E) of SJRHB031320_D1 following dataset integration. **F,** Bar graph comparing cellular diversity of single-cell and single-nucleus datasets of SJRHB031320_D1.

Extended Data Figure 2

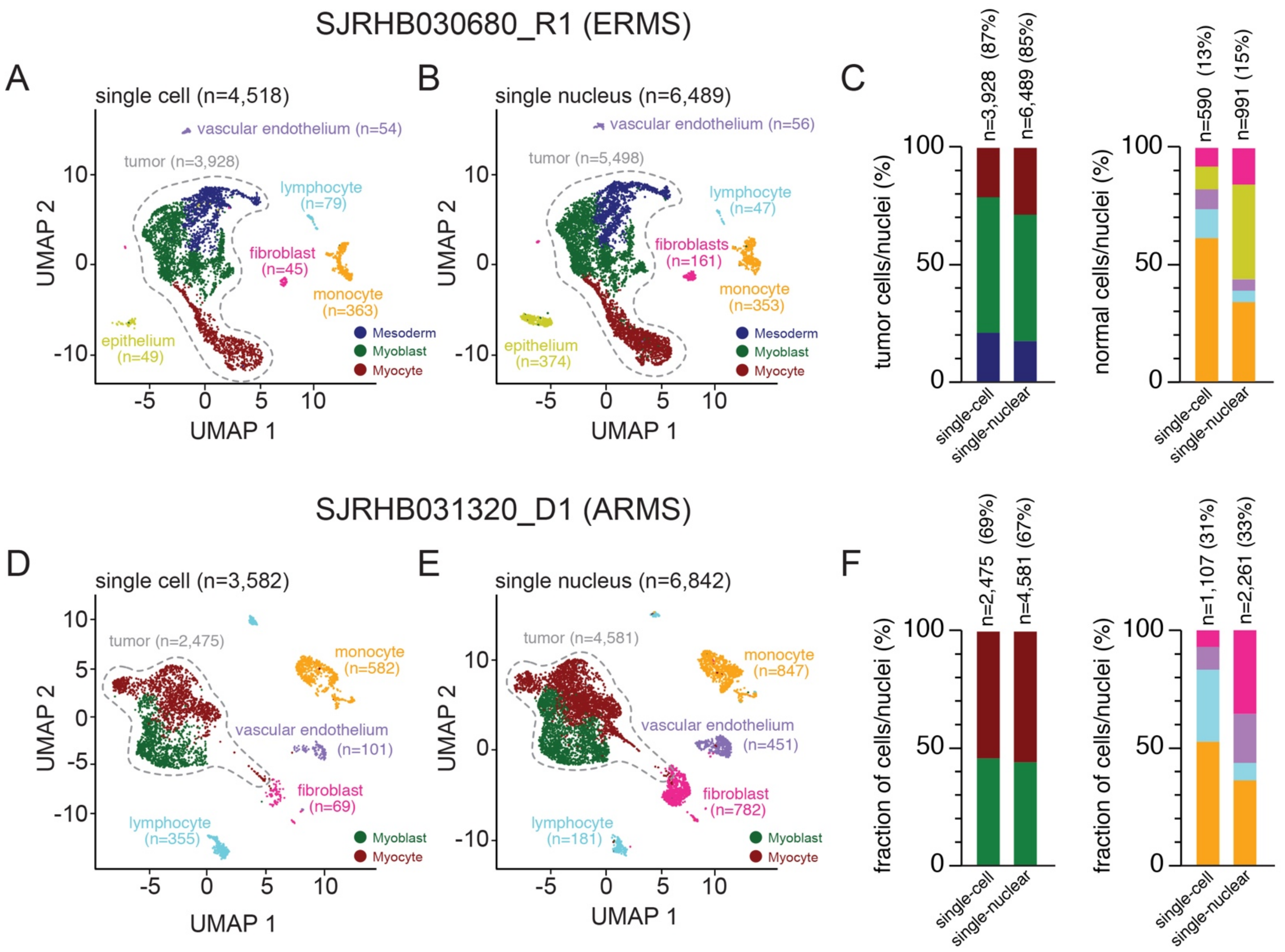

**Extended Data Figure 3: Combined analysis of malignant and non-malignant RMS nuclei.** **A,** Bar plots showing the percent composition of malignant nuclei within each developmental state. **B,** UMAP plots of SJRHB00026_R2, a relapsed ERMS sample obtained before treatment, and SJRHB010928_R1, an ERMS sample obtained after extensive treatment and noted to have <5% viable tumor cells on pathology. **C-D,** LargeVis plot of 11,357 non-malignant nuclei from snRNA-seq of patient tumors colored by sample (C) or by cluster (D). A heatmap showing expression of markers that were used for identification of cell types is shown on the right. **E,** Bar plots showing the percent composition of non-malignant nuclei within each sample. **F,** Plot showing fraction of non-malignant cells in each cell type cluster.

Extended Data Figure 3

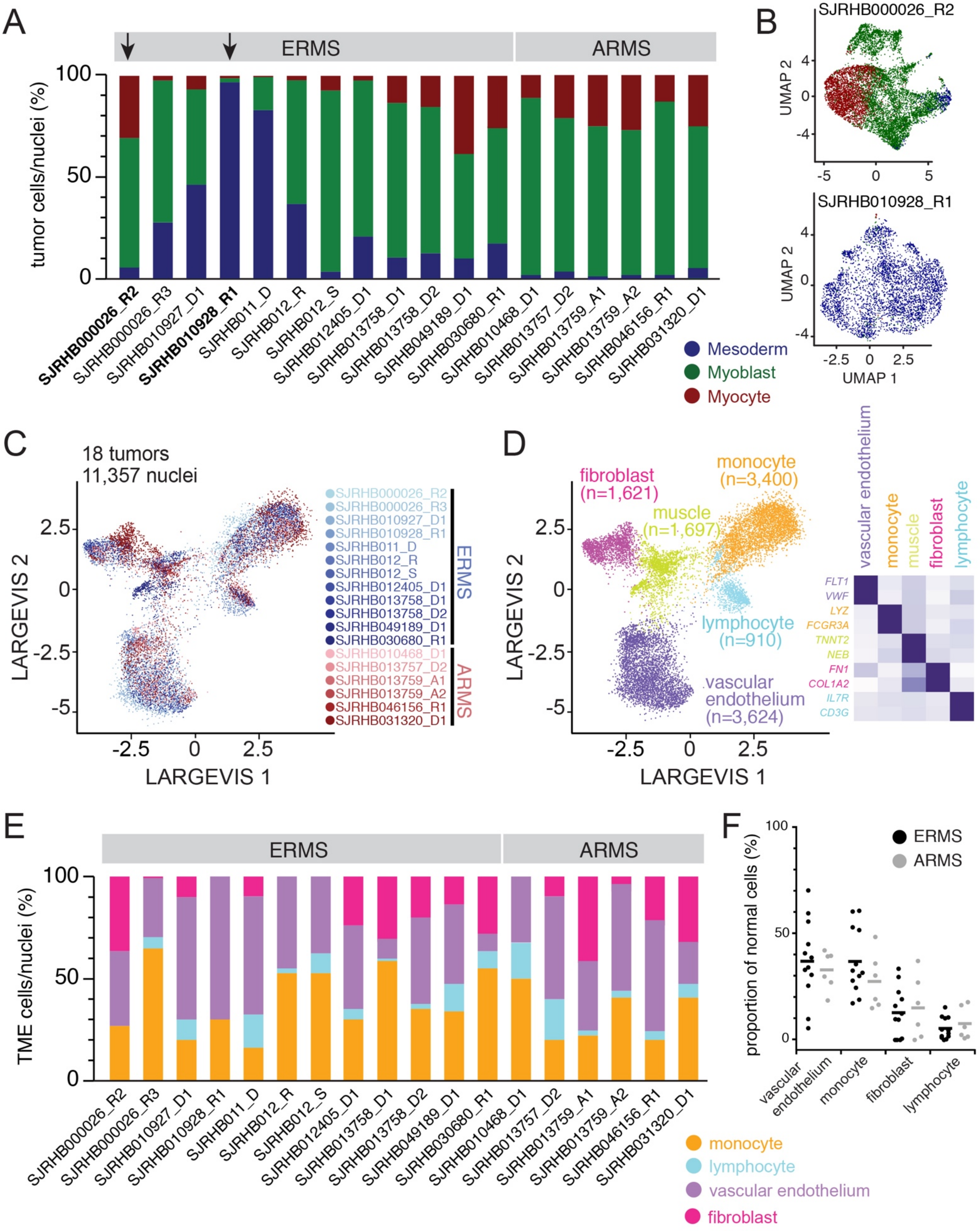

**Extended Data Figure 4: Combined analysis of single-cell RNA-seq data from RMS O-PDXs. A,** Bar plots showing the percent composition of malignant cells within each developmental state for 18 single-cell RNA-sequenced O-PDXs. **B,** UMAP plots of SJRHB010928_R1, an ERMS patient sample obtained after treatment that was almost entirely composed of cells in the mesoderm developmental state, and the tumor generated after xenotransplantation, SJRHB010928_X1, which has re-established the RMS developmental hierarchy. **C-E,** LargeVis visualization of scRNA-seq of 101,639 cells from 18 O-PDXs, coloured based on sample (C), Leiden cluster (D), or cell cycle phase (E). **F,** Comparison of single-cell/nucleus RNA-seq profiles from one patient tumor, SJRHB030680_R1 that underwent single-cell RNA-seq, single-nucleus RNA-seq, xenotransplantation, and xenotransplantation followed by ex vivo organoid formation. **G,** Quantitation of cell type proportions from panel F. **H,** Example IHC images of the O-PDX SJRHB011_X stained showing heterogeneous nuclear staining of MEOX2, MYF5, and MYOG. **I,** Quantitation of the percentage of cells within each O-PDX that are positive for MEOX2 (blue), MYF5 (green), or MYOG (red) immunohistochemical staining (*x* axis) compared to percentage of cells within each developmental state as determined by scRNA-seq (*y* axis).

Extended Data Figure 4

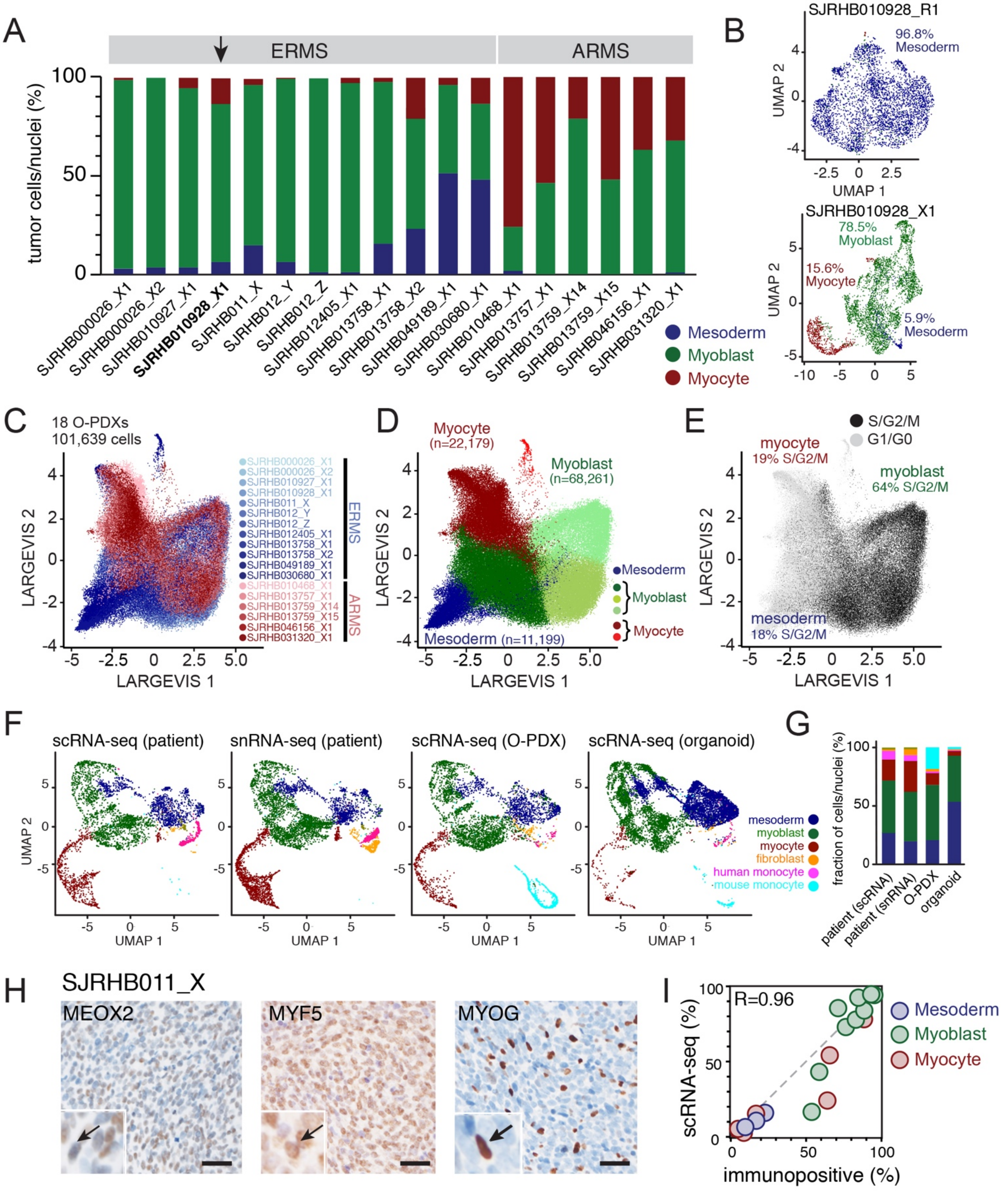

**Extended Data Figure 5: Representative CRC-SE in scATAC-seq. A,** scATAC-seq traces for the mesoderm, myoblast and myocyte populations for the *MEOX2* CRC-SE. **B,** violin plot showing expression of *MEOX2* in each population with enrichment in the mesoderm population. **C-E,** Peaks, CRC-SE boundaries and gene orientation for the *MEOX2* locus. **F,** scATAC-seq traces for the mesoderm, myoblast and myocyte populations for the *FOXO1* CRC-SE. **G,** violin plot showing expression of *FOXO1* in each population with enrichment in the mesoderm population. **H-J,** Peaks, CRC-SE boundaries and gene orientation for the *FOXO1* locus. Abbreviation: CRC-SE, core regulatory circuit super-enhancer.

Extended Data Figure 5

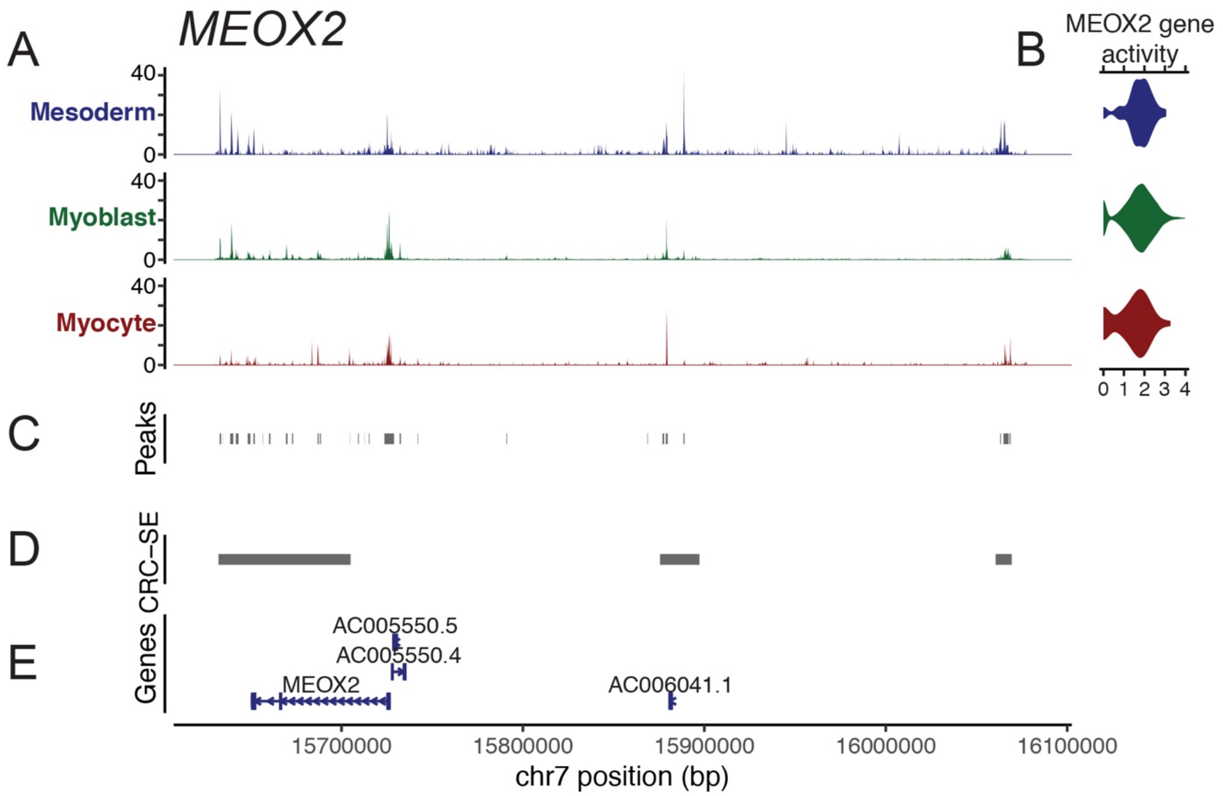


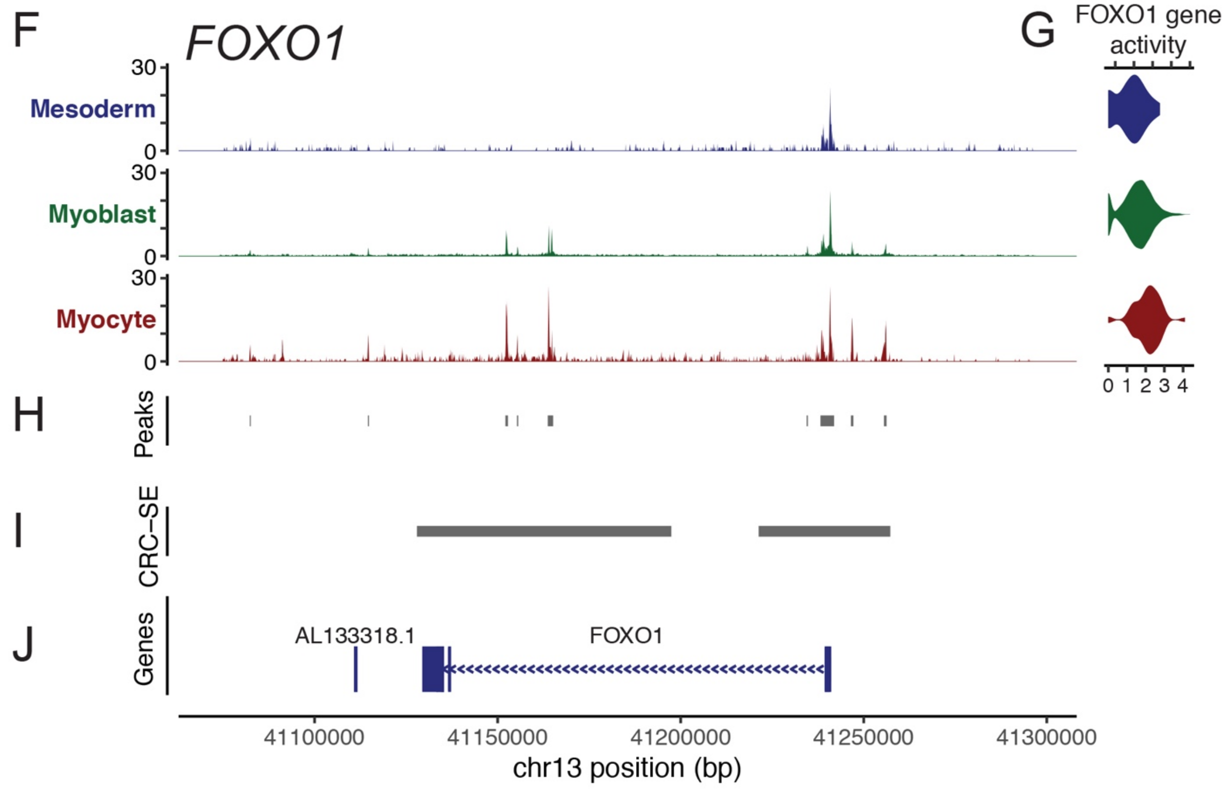

**Extended Data Figure 6: Targeting EGFR in 3D spheroids. A-C,** Representative IHC staining of an organoid generated from ERMS O-PDX, SJRHB010927_X1, stained for MEOX2 (A), MYF5 (B), or MYOG (C). **D-F,** Organoid viability, measured using CellTiter-Glo 3D, for SJRHB010927_X1 organoids treated with 3.9 nM, 15.6 nM or 250 nM of SN-38 for 72 hrs in the presence or absence of EGFR inhibitors (gefitinib or afatinib). Error bars, S.D of four technical replicates. **G,** Survival curves for each treatment group for a second ERMS tumor O-PDX SJRHB010927_X1.

Extended Data Figure 6

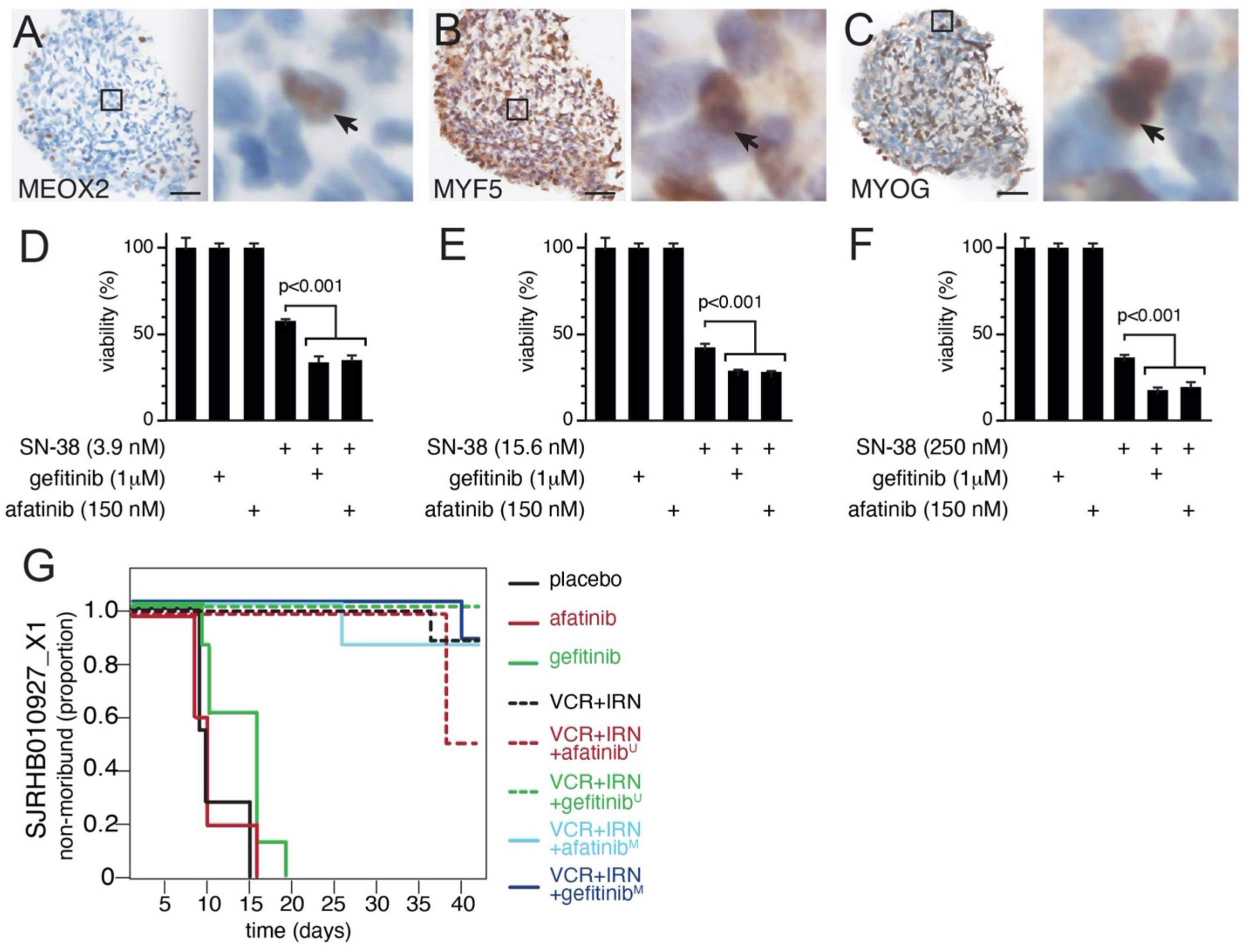

### Supplementary Materials and Methods

#### *Patient Samples*

Fresh primary patient rhabdomyosarcoma (RMS) tissue samples were obtained through the Molecular Analysis of Solid Tumor protocol (protocol ID XPD09-234) at St. Jude Children's Research Hospital[9]. All de-identified tissue samples were obtained after patient/family consent in agreement with local institutional ethics guidelines and institutional review board approval. Flash-frozen tissue samples were obtained through the St. Jude Children's Research Hospital Biorepository after approval by the St. Jude Children's Research Hospital Institutional Review Board (protocol ID XPD17-183). Formalin-fixed tissue from pre-treated and mid-treatment RMS resections were obtained as part of the RMS13 trial at St. Jude Children's Research Hospital (NCT01871766).

#### *Orthotopic Patient Derived Xenografts*

Orthotopic patient-derived xenografts (O-PDXs) described in this paper were obtained through the Childhood Solid Tumor Network (https://www.stjude.org/research/resources-data/childhood-solid-tumor-network.html)[9,31]. Clinical details related to these samples are provided in Supplementary Table 1. Single-cell suspensions of O-PDXs were diluted to 10,000 cells/µl in 100 µl Matrigel (BD Biosciences, catalogue number 354234) prior to intramuscular injection into the right hindlimb of mice. Female NSG mice (Jackson Laboratories, strain code 005557) were used for initial engraftment, and athymic nude mice (Charles River Laboratories, strain code 553) were used for passaging of O-PDXs. Tumor-bearing mice were euthanized once tumors reached 20% total body weight or once tumor size limited mobility.

To perform longitudinal biopsy sampling of O-PDXs, we anesthetized mice with inhaled 1-3% isoflurane. The skin overlying the tumor was sterilized with an alcohol prep before making a small, approximately 3 mm incision through the skin. A 23-gauge needle loaded with 0.3-0.5 cc of sterile saline was then passed through the tumor while applying negative pressure to the syringe plunger. The incision was then resealed using one drop of VetBond (3M). For pain control, mice were dosed with subcutaneous injections of 5 mg/kg Rimadyl (Zoetis) every 12 hours for 2 doses following the biopsy procedure.

Preclinical testing was performed in nude mice bearing luciferase-labelled O-PDXs as described previously[4,9]. After injection, mice were observed weekly and were randomly enrolled into treatment groups once tumors became large enough for a pretreatment biopsy (approximately 3 $mm^3$). Mouse enrollment in treatment groups are shown in Extended Data Tables 13 and 14. Vincristine was dosed at 0.19 mg/kg (50% dose group) or 0.38 mg/kg (100% dose group) once a week intraperitoneally, and irinotecan was administered at doses of either 1.56 mg/kg (50% dose group) or 3.125 mg/kg (100% dose group) on days 1-5; gefitinib and afatinib were orally gavaged daily at doses of 15 mg/kg and 30 mg/kg, respectively. Tumor volume was ascertained by Xenogen bioluminescence, and mice were euthanized on study once tumor volume exceeded 20% of total body weight. During chemotherapy treatment, mice were monitored daily.

#### *Tumor Dissociation*

*Fresh tissue*

Fresh tumor fragments from either patient or O-PDXs (< 500 mg) were rinsed with phosphate-buffered saline without calcium or magnesium (PBS-minus) prior to mincing with sterile scalpels. Enzymatic dissociation was then performed using components from the Papain Dissociation System (Worthington Biochemicals, catalog number LK003150). Tumor fragments were incubated in 5 ml papain-DNase solution at 37° C for 30 min (patient tumor) or 60 min (O-PDX tissue), followed by trituration with a 10 ml pipette and filtration through a 40 µm strainer. Single-cell suspensions were then pelleted at 500x*g* for 5 min. Cells were resuspended in resuspension buffer (2.7 ml Earle's buffered salt solution, 300 µl albumin-ovomucoid inhibitor solution, 150 µl DNase solution). Resuspended single-cell suspensions were layered over 5 ml albumin-ovomucoid inhibitor solution and centrifuged at 100x*g* for 6 min. The cell pellet was resuspended in PBS-minus buffer and filtered through a 40 µm strainer prior to downstream single-cell RNA-sequencing (scRNA-seq).

***Frozen tissue***

Flash-frozen tumor fragments were processed for single-nucleus RNA-sequencing (snRNA-seq) according to the TST extraction protocol described in Slyper et al[14]. Flash-frozen tumor fragments (approximately 50-100 mg) were incubated for 10 minutes in 1 ml TST buffer (73 mM sodium chloride, 5 mM Tris [pH 8.0], 0.5 mM calcium chloride, 10.5 mM magnesium chloride, 0.01% bovine serum albumin, 0.03% Tween-20) while mincing with Noyes spring scissors. Nuclei suspensions were then filtered through a 40 µm filter, followed by rinsing of the filter with an additional 1 ml of TST buffer. Nuclei suspensions were diluted with 3 ml of ST buffer (73 mM sodium chloride, 5 mM Tris [pH 8.0], 0.5 mM calcium chloride, 10.5 mM magnesium chloride) and centrifuged for 5 min at 500x*g* at 4° C. The nuclei pellet was then resuspended in 100-500 µl ST buffer and filtered through a 40 µm strainer prior to snRNA-seq.

For single-cell ATAC-sequencing (scATAC-seq), flash-frozen tumor fragments were processed according to the 10x Genomics recommended protocol for flash-frozen tissue with modifications. Frozen tumor fragments were placed in a microcentrifuge tube and lysed in 500 μl lysis buffer (10 mM Tris-HCl [pH 7.5], 10 mM sodium chloride, 3 mM magnesium chloride, 1% bovine serum albumin, 0.01% Tween-20, 0.01% NP-40, 0.001% digitonin). Tissue fragments were immediately homogenized 15 times using a microcentrifuge pellet pestle (Thermo Fisher Scientific). Homogenized nuclei suspensions were then incubated on ice for 10 min and diluted with 1 ml chilled wash buffer (10 mM Tris-HCl [pH 7.5], 10 mM sodium chloride, 3 mM magnesium chloride, 1% bovine serum albumin, 0.01% Tween-20). Nuclei suspensions were filtered through a 40 μm strainer prior to centrifugation at 500x*g* for 5 min at 4° C. Pelleted nuclei were resuspended in diluted nuclei buffer (10x Genomics), filtered through a 40 μm strainer, and then processed for scATAC-seq.

***Single cell/nucleus RNA-sequencing***

scRNA-seq and snRNA-seq were performed using version 2 or 3 of the 10x Genomics Single Cell RNA Expression Solution kits. Ten-thousand cells or nuclei were input into the 10x Chromium controller for droplet partitioning with barcoded beads. Barcoded libraries were generated according to manufacturer instructions. Each library underwent paired-end sequencing (50,000 paired end reads/cell) on an Illumina NovaSeq 6000 sequencer and processed using bclfastq to generate: Read 1 - 26 nucleotides, Read 2 - 100 nucleotides, Index - 8 nucleotides.

For barcoded O-PDXs, a separate dial-out PCR was performed on scRNA-seq libraries of barcoded O-PDXs using the primers 5’-TCGTCGGCAGCGTCAGATGTGTATAAGAGACAG

CTACACGACGCTCTTCCGAT-3' and 5'- GTCTCGTGGGCTCGGAGATGTGTATAAGAG ACAGTAGCAAACTGGGGCACAAGC-3'. Amplification was performed using Ex Taq polymerase (Takara) with 1 μl of scRNA-seq library and 80 nM of each primer. We used the PCR program:

98° C for 1 min

98° C for 10 sec, 65° C for 10 sec, 72° C for 20 sec for 35 cycles

72° C for 1 min

PCR product underwent clean-up using a PCR Purification Kit (Qiagen) or SPRIselect beads (Beckmann Coulter) before next-generation sequencing (5 million reads per sample, 100 bp paired-end reads) on an Illumina MiSeq sequencer. Sequencing data was then analyzed to generate cell identifier-barcode index tables using UMI-tools whitelist command[32]. The resulting index table was filtered to apply only those lineage barcodes for which at least 5 reads connected a barcode to a cell. The filtered table connecting 10x cell identifier to lineage barcode was then imported into Seurat metadata for downstream analysis.

#### *Single cell ATAC-sequencing*

scATAC-seq was performed using version 1.1.0 of the 10x Genomics Single Cell ATAC kit, according to manufacturer instructions. Two-thousand nuclei underwent transposition and were input into the 10x Chromium controller for barcoding, followed by library generation. Barcoded libraries underwent sequencing (50,000 paired-end reads/nucleus) on an Illumina NovaSeq 6000 sequencer and processed using bcl2fastq to generate: Read 1 – 50 nucleotides, Read 2 – 50 nucleotides, Index 1 – 8 nucleotides, Index 2 – 16 nucleotides.

***Data Analysis***

Fastq files were aligned to human hg19 genome for patient samples (10x Genomics; reference version 3.0.0) or combined hg19-and-mm10 genomes for O-PDX samples (10x Genomics; reference version 3.0.0) using the count command from Cell Ranger version 3.0.2 (10x Genomics). For samples generated by snRNA-seq, we included intronic counts to improve cell detection using a custom "pre-mRNA" genome reference[14]. The mkref command from Cell Ranger was utilised as described in https://support.10xgenomics.com/single-cell-gene-expression/software/pipelines/latest/advanced/references to generate the "pre-mRNA" reference.

Downstream analysis was performed in Seurat version 3.1.2[30,33]. Sparse matrixes were input from the Cell Ranger filtered_bc_matrix output and filtered to remove low quality cells and doublets. We excluded cells or nuclei with less than 400 genes, more than 7000 genes (presumed doublets), or cells where more than 10% of the unique molecular identifiers (UMIs) came from mitochondrial genes. Data was then normalised and transformed using scTransform version 0.2.1[34] within Seurat. Normalised, transformed data underwent principal component (PC) analysis of the 2000 most variably expressed genes. We used the top 30 PCs as input into Louvain algorithmic clustering with the resolution set to 0.4. Results were visualised by embedding cells or nuclei transcriptomic profiles using Uniform Manifold Approximation and Projection (UMAP)[35] of the top 30 PCs. Cell cycle stage prediction was performed using Seurat's built-in cell cycle scoring algorithm (https://satijalab.org/seurat/v3.0/cell_cycle_vignette.html) with cell-cycle markers identified by Tirosh et al[13]. To generate the marker UMAP plots in Fig. 1e and 1f, cells are colors based on the level of expression of the listed marker (MEOX2, MYF5, or MYOG). Marker-positive fractions were calculated using the number of cells with $\log_2$ normalized expression > 1 (“marker-positive cells”) divided by the total number of cells in the dataset. To perform RNA velocity analysis of SJRHB030680_R1 and SJRHB031320_D1, the post-sorted bam files from the Cell Ranger count pipeline was input into velocyto version 0.17.17[15], and analyzed using Scanpy version 1.4.5[36] and scVelo version 0.1.24be[37].

For each cell subset identified by clustering, we used a combination of SingleR version 1.0.1[38] and manual inspection of differentially expressed genes to annotate whether a cluster belongs to stromal, immune or malignant subpopulations. Malignant cells were confirmed in patient tumor data by inference of copy-number variation using inferCNV version 1.1.3 of the TrinityCTAT Project (https://github.com/broadinstitute/infercnv). An average read count per gene cutoff of 0.1 was used. Reference cells were defined by those cell clusters in the Seurat analysis that expressed markers of hematopoietic or endothelial cells. Analysis included denoising, implementation of inferCNV’s i6 HMM model, and subclustering using quantile normalization.

For the comparison of tumors that underwent both scRNA-seq and snRNA-seq in Extended Data Fig. 2 and Fig. 5K, we utilized Seurat’s integration workflow to identify anchors and integrate scRNA-seq and snRNA-seq datasets across the first 30 canonical components[30]. For the large-scale integration of 18 snRNA-seq datasets from patient RMS tumors (Fig. 2), we used Conos version 1.2.1[16] to build a unified graph of the datasets (buildgraph settings: k=30, k.self=5, n.odgenes=2000, alignment.strength=0), with mutual nearest-neighbor mapping of the first 50 common PCs. Shared community clustering was performed using the Leiden algorithm[39] with resolution 0.4, followed by UMAP visualization. Wilcoxon Rank Sum testing was performed within Seurat using the FindAllMarkers command to identify differentially expressed genes.

Mesenchymal developmental snRNA-seq data from previously published mouse organogenesis data[18] was downloaded from https://oncoscape.v3.sttrcancer.org/atlas.gs.washington.edu.mouse.rna/landing, imported into Seurat, and clustered with resolution 0.8 to identify subclusters associated with muscle development (Extended Fig. 7). Those subclusters were extracted to generate the mouse skeletal muscle development dataset used for downstream PC projection and trajectory analysis. Trajectory inference was performed on mouse skeletal muscle development snRNA-seq data using Slingshot via the dynverse platform[40,41]. Output from Slingshot underwent lineage and pseudotime assignation.

scATAC-seq data were processed using the Cell Ranger ATAC version 1.2.0 (10x Genomics). Cell Ranger ATAC output was then imported into R via Signac version 1.1.0 (https://github.com/timoast/signac). Data was filtered to evaluate only those cells with greater than 20% of reads within peak fragments, between 1000 and 20000 unique fragments within peak regions, less than 5% of peaks within ENCODE-defined blacklisted regions, a nucleosome signal (defined as the ratio of mononucleosome fragments to nucleosome-free fragments) less than 10%, and greater than two-fold enrichment at transcription start sites as defined by ENCODE. Filtered datasets underwent term frequency-inverse document frequency normalization, followed by singular value decomposition for dimensionality reduction. We used the top 20 dimensions for UMAP non-linear dimensional reduction, visualization, and graph-based clustering. To integrate scATAC-seq and scRNA-seq data, we estimated gene activities in the scATAC data using the "GeneActivity" command of Seurat, which was used to perform cross-modality and label transfer within Seurat. Transferred cluster assignment were input into Loupe ATAC (10x Genomics) to generate subpopulation-specific chromatin peak accessibility profiles. For detection of peaks within previously determine CRC-SE regions[4], we generated feature matrices in Signac using the defined CRC-SE peak ranges; after normalization, we performed differential accessibility analysis using the FindMarkers command in Seurat using logistic regression while using the total number fragments as a latent variable for analysis.

#### *Developmental Indexing*

Latent cellular states identification: Mesenchymal developmental snRNA-seq data was combined with snRNA-seq data from SJRHB030680_R1 (ERMS) and SJRHB031320_D1(ARMS) for latent cellular states (LCs) identification[19]. Briefly, the general difference between malignant (two RMS tumors) and normal mouse developmental muscle cells were corrected. LCA clustering analysis of the developmental muscle data only revealed both a similar structure with the subpopulations identified in Seurat (mesoderm, paraxial mesoderm, myoblast, myocytes and myotubes) and a set of 20 LCs that supports the distinguishing of the subpopulations.

Muscle developmental index projection: LCs for individual cells were derived from their global expression profiles[19]. The similarity between a testing cell *b* to a training cell *a* is calculated as:

$$similarity_{a,b} = \frac{\sum_{i=1}^{p} LC_{a,i} \times LC_{b,i}}{\sqrt{\sum_{i=1}^{p} {LC_{a,i}}^2} \times \sqrt{\sum_{i=1}^{p} {LC_{b,i}}^2}}$$

where $p$ represents the number of LCs retained in the previous step (20 in this analysis). The normalized similarity for testing cell $b$ is defined as:

$$normalized\_similarity_{a,b} = \frac{similarity_{a,b}}{sd(similarity_{.,b})}$$

where $similarity_{.,b}$ represents similarity scores between testing cell $b$ and all cells in training datasets. The raw developmental index was derived from a weighted average of the $k$ (default 25) nearest neighbor cells in the training samples:

$$Developmental\_index_b = \frac{\sum_{a=1}^{k} w_{a,b} \times PS_a}{\sum_{a=1}^{k} w_{a,b}}$$

where $PS_a$ represents the Slingshot inferred pseudo-temporal output for cell $a$ in the training data and $w_{a,b}$ represents the weight between testing cell $b$ and one of the nearest neighbor cell $a$ in the training data:

$$w_{a,b} = e^{\frac{normalized\_similarity_{a,b}}{2}}$$

Finally, the developmental index was normalized using the empirical cumulative distribution function estimated from the training dataset:

$$Normalized\_developmental\_index_b = \frac{\sum_{a=1}^{n} I(PS_a \leq Developmental\_index_b)}{n}$$

where $n$ represents the number of cells in the training dataset and $I(true/false) = 1/0$, respectively.

***NetBID Analysis and driver identification from bulk RNA-seq data and from snRNA-seq***

The NetBID (data-driven network-based Bayesian inference of drivers) algorithm[23,24] was used to identify drivers in ERMS and ARMS patients from bulk RNA-seq profiles of our published RMS cohort[4]. An RMS-specific signaling interactome (RMSi) from 77 RNA-seq profiles of

RMS patients was generated using SJARACNe[25], an information theory-based algorithm for regulatory network inference. The parameters of SJARACNe were configured as the following: p value threshold p = 1e-7, data processing inequality (DPI) tolerance e = 0, and number of bootstraps (NB) = 100. After generating the RMSi, the "weighted mean" algorithm (*cal.Activity* function) in NetBID was utilized to infer activities of signaling driver candidates (e.g., EGFR) across 52 ERMS and 25 ARMS patient samples from their gene expression profiles. Drivers in ERMS and ARMS were identified using the *getDE.BID.2G* function in NetBID.

To adapt NetBID to snRNA-seq data, the SJARACNe algorithm was used to computationally reconstruct cell type–specific interactomes for cell types that have >2,000 nuclei from the integrated snRNA-seq profiles. It resulted in 5 signaling networks for cell types of ERMSmesoderm (19,317 nuclei, 11,215 genes, 195,707 edges), ERMSmyoblast (43,667 nuclei, 11,182 genes, 257,252 edges), ERMSmyocyte (10,674 nuclei, 11,221 genes, 732,180 edges), ARMSmyoblast (29,040 nuclei, 12,374 genes, 122,389 edges), and ARMSmyocyte (7,757 nuclei, 11,222 genes, 331,461 edges). With a focus on signaling proteins, the cell type–specific interactomes of 2,543 signaling genes were used to infer network activities in each nucleus using the interactome of the corresponding cell type. To overcome the sparseness of snRNA-seq data, the "unweighted mean" algorithm in the *cal.Activity* function of NetBID was used. Differential activity analyses to identify cell type–specific drivers were performed by using the *getDE.BID.2G* function in NetBID.

***Real-time PCR via Taqman Array Cards***

Real-time PCR experiments were performed using the Applied Biosystems QuantStudio Flex 7 instrument. O-PDX tissue obtained after dissection or biopsy underwent RNA extraction using Trizol reagent (Invitrogen) as per manufacturer instructions. One microgram of extracted RNA was used for cDNA synthesis using the High Capacity RNA-to-DNA kit (Invitrogen). cDNA libraries were then mixed with TaqMan Fast Advanced Master Mix (Invitrogen) and loaded onto custom TaqMan Array Cards (Invitrogen) using primers directed against 23 human genes listed in Extended Data Table 8-12. Samples were analyzed in duplicate, using *GAPDH* levels for normalization. To generate mesoderm signature scores, we averaged the fold-changes of *MEOX2*, *PAX3*, *EGFR, DCN, CD44* and *POSTN*; for myoblast signature scores, we averaged the fold-change of *PAX7*, *MYF5, MSC, GPC3,* and *VIM*; for myocyte signature scores, we averaged the fold-change of *MYOD1, MYOG, MEF2A, MEF2C, TTN, NCAM1, MYH3, NEB,* and *CDH15*.

***Immunohistochemistry***

All formalin-fixed, paraffin-embedded (FFPE) tissues were sectioned at 4-μm, mounted on positively charged glass slides (Superfrost Plus; Thermo Fisher Scientific, Waltham, MA), and dried at 60° C for 20 min. Procedures and antibodies used to detect human antigens to MEOX2, MYF5, MYOG and EGFR are listed below:

| **Antibody** | **Type** | **Concentration** | **Supplier, Catalog Number** |
|---|---|---|---|
| EGFR | Rabbit monoclonal (5B7) | RTU[a] | Ventana/Roche, 790-2988[b] |
| MEOX2 | Rabbit polyclonal | 1:500 | Sigma Aldrich, HPA053793[b] |
| MYF5 | Mouse monoclonal (OTI2G5) | 1:200 | Invitrogen, MA5-26654[c] |
| MYOG | Mouse monoclonal (F5D) | 1:150 | Agilent, M3559[d] |

[a]RTU: ready to use

[b]Ventana DISCOVERY ULTRA automated stainer (Ventana Medical Systems, Inc., Tucson, AZ): Heat-induced epitope retrieval (HIER), Cell conditioning media 1 (CC1), 32 minutes at 37 °C; Visualization with DISCOVERY OmniMap anti-rabbit HRP (760-4311), DISCOVERY ChromoMap DAB kit (760-159) or DISCOVERY ChromoMap Purple kit (760-229)

[c]Leica BOND-MAX automated stainer (Leica Biosytems, Buffalo Grove, IL): IHC Protocol F. HIER with Bond Epitope Retrieval Solution 2 (ER2) for 20 minutes; Visualization with Bond Polymer Refine Red Detection (DS9390).
[d]Ventana DISCOVERY ULTRA automated stainer (Ventana Medical Systems, Inc.): HIER, Cell conditioning media 2 (CC1), 32 minutes at 37 °C; Visualization with biotinylated rabbit anti-mouse IgG secondary antibody (Abcam, ab133469), DISCOVERY OmniMap anti-rabbit HRP (760-4311), DISCOVERY ChromoMap DAB kit (760-159) or DISCOVERY ChromoMap Purple kit (760-229)

Human RMS O-PDX samples that were sequenced and shown to express the protein targets at the mRNA level were used as positive tissue controls for IHC. A neuroblastoma O-PDX sample was used as a negative tissue control. Isotype controls were used for monoclonal antibodies where appropriate.

***Lentiviral Barcoding***

pBA439 barcode library[20,21] (Addgene, catalog #85968) was expanded in ElectroMAX Stbl4 *E. coli* cells (Thermo Fischer Scientific) by electroporating 50 ng of barcode library into 100 µl *E. coli* using a BioRad GenePulser II instrument with the settings: 1.2 kV, 25 µF, 200 Ω. Electroporated cells were expanded in 4 L Luria broth supplemented with 100 µg/ml carbenicillin (Sigma Aldrich) for 16 hr at 37°C with shaking before isolation of plasmid DNA via Maxiprep Plasmid Kits (Qiagen).

Lentivirus was generated using ten 100mm tissue culture dishes of 40% confluent HEK293T cells. Each plate of cells underwent transfection of 6 µg of the pBA439 plasmid library along with third-generation lentiviral packaging vectors[42] (3 µg CAG-kGP1.1R, 1 µg CAG4 RTR2, and 1 µg HDM-G) using polyethyleneimine (Sigma-Aldrich) at a ratio of 1:2 DNA:polyethyleneimine. Viral supernatant was collected at 48 and 72 hr, followed by filtration through a 0.45 µm cellulose acetate filter (Corning). Viral particles were then concentrated by layering over a 20% sucrose gradient and ultracentrifuged at 24,000 rpm for 90 min at 4°C within a Beckman SW 32 Ti rotor. Concentrated viral particles were stored at -80°C, and titred to calculate transduction efficiency.

To generated barcoded xenografts, single-cell suspensions of O-PDXs were transduced by incubating 5 million cells with lentivirus ($1x10^6$ transducing units) in 1 ml DMEM media supplemented with 8 μg/mL polybrene (Sigma-Aldrich). Cells were incubated for 2 hr at room temperature, pelleted at 500x*g* for 5 min, and washed once with sterile DMEM. Washed, transduced cells were injected orthotopically into the hindlimbs of female nude mice and allowed to grow to approximately 20% animal body weight. Post-transduction xenografts were dissected, dissociated and sorted on a S3e cell sorter (Bio-Rad) for blue fluorescent protein (BFP)-positivity and 7-aminoactinomycin D (Invitrogen) exclusion to select for viable, barcoded cells. Sorted, barcoded cells were re-injected into female nude mice diluted in 0.1 ml Matrigel at 1,000 cell/μl to generate barcode-enriched O-PDXs.

***Mathematical modeling***

We simulated tumor growth in vivo by using a 3-dimensional agent-based model that follows the fate of cells in both space and time. It contains three populations: mesoderm, myoblast, and myocytes. Basic cell division, differentiation, and death processes are modeled. During each time step, mesoderm cells divide on average with a probability $L_{mes}$. With a probability $P_{mes}$, a self-renewal division occurs, generating 2 mesoderm daughter cells. With a probability $1-P_{mes}$, a differentiation division occurs, generating two myoblast cells. We assumed that self-renewal divisions are more likely than differentiating divisions ($P_{mes}>0.5$), i.e. the mesoderm population expands over time. Myoblasts followed similar dynamics. They were modeled to divide with a probability $L_{blast}$. This division results in self-renewal with a probability $P_{blast}$ (creating two myoblast cells), and in differentiation with a probability $1-P_{blast}$ (creating two myocyte cells). We assumed that myoblasts on their own cannot sustain growth ($P_{blast} < 0.5$). Myocytes were assumed not to divide, and they die with a probability D. When a division event occurs, one of the 27 neighboring locations is chosen randomly. The offspring is placed there if this spot is empty, otherwise the division event is unsuccessful, due to density dependence. To account for the observation that *47%* of mesoderm cells were in a dividing state, we assumed that a certain fraction of the cells, *f*, are dividing faster, while the remaining cells were assumed to be more quiescent and divide only infrequently, such that the average division probability across all mesoderm cells is $L_{mes}$ ($f=(1-r_2)/(r_1-r_2)$, where $r_1$ and $r_2$ are the dimensionless slow and fast division rates, respectively). In accordance with data, we further assumed that myoblast cells divided with a rate that was 1.3 times faster than the average rate of mesoderm divisions. The simulated tumors initially consisted of 1.3% mesoderm cells, 97.4% myoblast cells, and 1.3% myocytes, based on the initial experimental conditions. The mesoderm and myoblast compartments were seeded probabilistically with 33 bar codes, according to the experimentally documented distribution. The fate of the individual barcodes was tracked over time.

Treatment was simulated by including the death of dividing cells. Thus, all myoblast cells were assumed to die with a relatively fast rate $\alpha_{blast}$; dividing mesoderm cells were assumed to die with a slower rate $\alpha_{mes}$, while quiescent mesoderm cells were assumed to be resistant against treatment-induced death. Upon treatment cessation, we assumed a reactivation of a certain portion of remaining quiescent mesoderm cells, to be consistent with the experimentally observed fraction of mesoderm cells in G2/M.

The “entropy index” is a number that measures the diversity of different populations. The lower the entropy index, the more uneven the distribution of bar codes among the cells, indicating dominance by only a few bar codes. Denoting the fraction of each bar code in the population by $x_i$, the entropy is given by $E = -\frac{x_i}{\sum x_i} ln \frac{x_i}{\sum x_i}$.

***RMS Organoids***

Single-cell suspensions of O-PDX were generated, as described previously[9], were washed with DMEM media (Thermo Fischer Scientific) and pelleted at 500x*g* for 5 min. Cells were resuspended to a concentration of 500,000 cells/ml in SkBM-2 skeletal muscle cell growth basal medium supplemented with muscle growth SingleQuot supplements (Lonza). 100 µl of cell suspension were aliquoted into wells of ultra-low attachment round-bottom Lipidure coated 96-well plates (Gel Company, catalogue number LCU96). Plates were spun at 300x*g* for 3 min to aggregate cells and allowed to grow incubated at 37° C. SkBM-2 media was exchanged weekly during organoid culture.

For organoid viability studies, organoids were grown 14 days before exposure to combinations of SN-38 with or with EGFR inhibitors (150 nM afatinib or 1 µm gefitinib) for 72 hrs. Cell viability was measured by adding 100 µL of CellTiter-Glo 3D (Promega catalog number G9681) to each well, followed by gentle agitation for 30 min.  The plate was read on a Pherastar plate reader (BMG LabTech).

***Data and Source Code Availability***

NetBID: https://jyyulab.github.io/NetBID

All processed single-cell and single-nucleus RNA-sequencing data are publicly accessible via an online data portal (https://pecan.stjude.cloud/static/RMS-scrna-atlas-2020).
